# Sex-specific gene and isoform expression characterizes hippocampal aging in rhesus macaques

**DOI:** 10.64898/2026.11.04.764396

**Authors:** Victoria Harper-Alexander, Sam Peterson, Saida Sadikova, Steven Kohama, Daniel Ramos Ram

## Abstract

Aging produces widespread transcriptomic alterations that extend beyond changes in gene expression to include changes at the isoform level. Declining splicing fidelity, including increased intron retention and aberrant isoform expression, has been proposed to contribute to cellular dysfunction; however, the extent and biological significance of isoform-specific changes in the aging brain remain poorly understood. Age-related cognitive decline has also been associated with neuroinflammation and the development of a chronic inflammatory state, or “inflammaging,” within the brain. Here, we used short- and long-read RNA sequencing to characterize gene- and isoform-level changes associated with hippocampal aging across sex in 24 rhesus macaques. We identified 594 age-associated differential isoform expression events, comprising both shared and sex-specific changes. Inflammatory pathways, including TNF-α–NF-κB signaling, interferon responses, and complement pathways, were enriched across both sexes. Weighted gene co-expression network analysis (WGCNA) further identified sex-specific modules associated with inflammatory, glial, mitochondrial, and hormone-responsive processes. Notably, several age-associated changes were detected at the isoform level despite limited or absent corresponding gene-level differences, highlighting transcript-level regulation as an additional dimension of brain aging that may be overlooked by gene-level analyses. To investigate the relationship between aging and Alzheimer’s disease (AD), we independently performed WGCNA on the rhesus macaque cohort and a publicly available human hippocampal AD single-nucleus RNA-sequencing dataset (GSE268609). Cross-species module preservation and gene overlap analyses identified shared genes between age-associated and AD-associated networks, with pathway enrichment implicating glial function, inflammation, and mitochondrial biology. Together, these findings highlight sex-associated differences in hippocampal transcriptomic aging, underscore the importance of isoform-level regulation, and identify shared transcriptional signatures between normal aging and AD-associated pathways.

## Introduction

Aging is a complex biological process driven by interconnected cellular and molecular processes that occur throughout an organism’s lifespan^1^. Our rapidly aging population is driving a rise in age-associated diseases, placing substantial pressure on healthcare systems worldwide^2^. Among these conditions, neurodegenerative diseases including Alzheimer’s disease (AD), Parkinson’s disease (PD), and related dementias account for a major portion of morbidity in older adults^3,4^. These disorders increase sharply with age and are characterized by progressive loss of neuronal and synaptic integrity, ultimately resulting in cognitive decline, functional impairment, and reduced quality of life. While individual genetic variants contribute to risk^5, 6^, most cases lack a single causal mutation, indicating that the process of aging itself is a primary factor that shapes vulnerability to neurodegeneration^7^. In the brain, these age-associated changes can manifest as a progressive neuroinflammatory landscape characterized by persistent glial activation^8^, impaired inflammatory resolution, and the accumulation of pathological protein aggregates such as β-amyloid, α-synuclein, and hyperphosphorylated tau^9^. Age-related immune dysregulation culminates in “inflammaging”^7^, a pervasive and chronic inflammatory state that spans peripheral tissues and the central nervous system. This systemic inflammatory milieu is linked to widespread physiological deterioration, including synaptic dysfunction^10^, neurodegeneration^7^, and cognitive decline^11^. Importantly, emerging data reveal that inflammaging is sexually dimorphic^12^ which adds another layer of complexity. Because of the heightened risk of neurodegenerative disorders, uncovering the regulatory mechanisms that connect normal aging to disease susceptibility is essential for identifying the pathways that dictate disease vulnerability. Transcriptomic profiling has revealed that aging remodels inflammatory networks in the brain^13^, but most studies rely on short-read RNA sequencing, which cannot fully capture additional complexities that influence gene expression such as alternative splicing (AS).

Alternative splicing is a core mechanism of tissue-specific gene regulation, that occurs in more than 90% of genes containing multiple exons^14^. The diversity of RNA transcripts reflects the interplay of multiple regulatory layers, including alternative transcription initiation (transcription start site [TSS] usage), alternative splicing (AS), and alternative polyadenylation (APA)^15^. In the central nervous system, AS is particularly prevalent and tightly regulated, playing essential roles in neuronal differentiation, synaptic plasticity, and cell-type specific communication^16^. Dysregulated AS is implicated in multiple age-related neurodegenerative diseases, including AD, PD, and frontotemporal dementia. For instance, mis-splicing of MAPT exon 10 disrupts 3R/4R tau homeostasis and promotes AD pathogenic aggregation, while altered SNCA isoform profiles contribute to the neurodegenerative pathology of Parkinson’s disease and frontotemporal dementia^17,18^. These examples illustrate that neurodegeneration is not solely driven by changes in gene-level abundance, but also by shifts in isoform composition. Emerging evidence suggests that aging may be associated with AS changes^19^. Instead of producing uniform increases or decreases in transcript abundance, aging can shift isoform usage toward non-canonical transcripts^20,21^. This shift, termed isoform switching, alters the balance of functional transcript variants. Isoform expression can fundamentally alter protein structure, signaling capacity, and cellular responses without changes in gene-level expression^22,23^. Importantly, many genes central to immune signaling and inflammatory regulation are regulated by alternative splicing^24, 25^, suggesting that age-associated isoform switching may represent a key mechanism by which aging reshapes inflammatory pathways.

Such regulatory remodeling is likely to be particularly consequential in brain regions that are highly sensitive to aging. The hippocampus, is a deep brain structure central to learning and memory and is among the earliest to exhibit age-related functional decline and neurodegenerative pathology^26,27,28^. With advancing age, the hippocampus develops a pronounced inflammatory milieu characterized by sustained glial activation and enrichment of immune-related signaling pathways^29^. Elevated hippocampal inflammation is also a defining feature of AD and other neurodegenerative diseases^30^, where it contributes to synaptic dysfunction and neuronal loss^31^. Despite the central role of inflammation in hippocampal aging, how alternative splicing and isoform switching regulate inflammatory signaling in this region remains poorly understood. Together, these observations suggest that aging-associated changes in AS may reprogram inflammatory pathways at the isoform level, thereby contributing to hippocampal vulnerability and increasing susceptibility to neurodegenerative disease. Defining how isoform-level regulation intersects with inflammation in the aging hippocampus is therefore critical for understanding the molecular mechanisms that link normal aging to neurodegeneration. Also, given that biological sex significantly impacts neurodegenerative disease succeptability^32, 33^ and inflammaging^34^, it is also necessary to understand how sex influences molecular changes in the brain at the level of AS. Rhesus macaques (Macaca mulatta) provide a powerful, translational model for addressing this gap due to their conserved neuroanatomy^35^, similar life stages and age-related hormonal progression^36^, shared immune architecture^37^, and genetic and physiological similarities to humans^38^. Importantly, rhesus macaques also exhibit age-associated neurodegenerative neuropathologies^39, 40^. While a recent large-scale short-read transcriptomic study of 96 rhesus macaque hippocampi identified 2,679 age-associated differentially expressed genes, no sex differences in the aging transcriptome were detected^41^. These findings raise the possibility that sex-specific effects of aging may be more apparent at the transcript isoform level than at the gene level. Building on this work, utilized Oxford Nanopore long-read RNA sequencing in a subcohort of 24 animals from the original study and leveraged the corresponding Illumina short-read RNA sequencing data from these same animals to characterize gene-level transcriptional changes. Integrating these complementary sequencing approaches, we investigated age- and sex-associated changes in gene and transcript isoform expression and examined whether age-associated transcriptional changes were enriched in genes and pathways implicated in neurodegenerative disease and inflammation.

## Results

### Study design and sequencing quality assessment

To characterize age-related transcriptomic alterations at isoform-level resolution, we performed long-read RNA sequencing on hippocampal tissue from 24 rhesus macaques (12 males and 12 females) (Fig. 1a). 22 rhesus macaques were reliably demultiplexed in the long-read sequencing assays and were used for further analyses. Within each sex, six individuals were classified as young (mean age = 5.42 years; males = 5.12 +/− 0.4 SEM, females = 5.72 +/− 0.36 SEM) and five as aged (mean age = 25.60 years; males = 25.46 +/− 0.65 SEM, females = 25.73 +/− 0.82 SEM) (Fig. 1b). Mean age was comparable between males and females within both the young and aged cohorts. Long-read sequencing was employed to overcome the inherent limitations of short-read RNA-seq, which fragments RNA molecules and may introduce gaps in transcript coverage (Fig. 1c), leading to inaccurate quantification and incomplete identification of alternatively spliced isoforms. Following basecalling, quality control, and alignment, gene-level and transcript-level expression matrices were generated for downstream differential expression and splicing analyses (see Methods and Supplementary Tables 1-2). Principal component analysis (PCA) of normalized transcript expression values revealed segregation of samples by both age and sex (Fig. 1d), indicating global transcriptomic differences associated with aging. Read quality was assessed using ToulligQC as previously described^42^. Within samples, read lengths ranged from 300-1800bp (Fig. 1e). Further read length and quality scores distributions are provided in Supplementary Figure 1 a-h. Together, these results outline the quality of the sequencing dataset, establishing a robust foundation for identifying isoform-level changes associated with hippocampal aging.

**Fig. 1:**
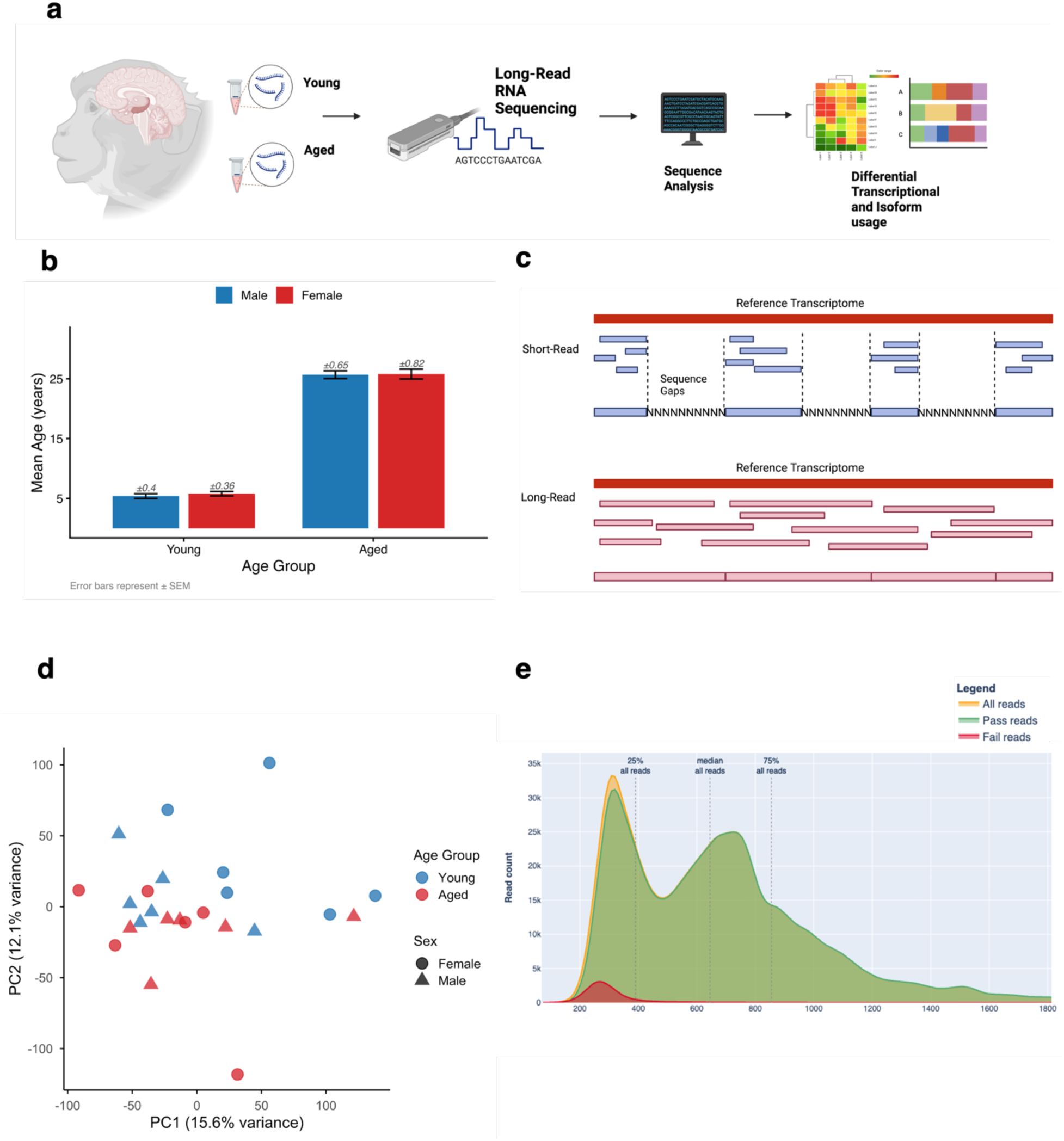
Experimental design and sample overview. **a** Overview of the experimental design showing collection of hippocampal tissue from young and very aged rhesus macaques for long-read RNA sequencing. **b** Mean age of young and aged groups; error bars represent +/− SEM. **c** Schematic comparison of short-read and long-read sequencing approaches; short-read sequencing introduces coverage gaps that limit isoform-level resolution, whereas long-read sequencing enables full-length transcript characterization. **d** Principal component analysis (PCA) of variance-stabilized transcript expression showing sample clustering by age and sex; PC1 and PC2 explain 15.6% and 12.1% of total variance, respectively. **e** Read length distribution plot of all reads passed and failed.

### Male hippocampal aging is associated with inflammatory and lipid metabolic transcriptional changes

To get a sense of transcriptomic changes with age at the gene level, we performed differential gene expression analysis using DESeq2. Differential expression testing was performed separately within each sex using the Wald test (Benjamini–Hochberg FDR < 0.05). This yielded 44 significant differientally expressed genes (DEGs) unique to females, 156 unique to males, and 8 DEGs shared across sexes. (Fig. 2a). A detailed analysis protocol is outlined in the methods section. In males, age-associated differential expression revealed a broad inflammatory and myeloid-associated transcriptional program (Fig. 2b). *TREM1*, an amplifier of innate immune and myeloid inflammatory signaling, showed a male-specific increase (log₂FC = +4.31, padj = 1.6 × 10⁻⁶), accompanied by increased expression of *IL1B* (+1.75, padj = 0.030), *CD68* (+1.27, padj = 0.041), *SPP1* (+0.94, padj = 0.038) and the acute-phase serpin genes *SERPINA3* (+2.72, padj = 0.014) and *SERPINA1* (+1.18). Additional increases in *NCF1* (+2.07, padj = 0.020), *RAC2* (+2.80, padj = 0.030), *GPR84* (+4.39, padj = 0.016), *CXCR4* (+1.09, padj = 0.014) and *PILRA* (+2.11, padj = 0.029) further supported coordinated remodeling of myeloid-associated genes (Supplementary Table 3). Male aging was also associated with increased expression of lipid- and lipoprotein-associated genes, including *APOD* (+0.83), *APOC2* (+2.43) and *LCAT* (+1.25), together with increased *TF* (+0.84), a mediator of iron transport. Astroglial- and extracellular matrix-associated transcripts, including *SPARC* (+0.79, padj = 3.5 × 10⁻⁵), *GPR37L1* (+0.51) and *FAM107A* (+0.46), were similarly increased, extending the age-associated remodeling beyond inflammatory myeloid programs to glial and tissue-support processes.

**Fig. 2.**
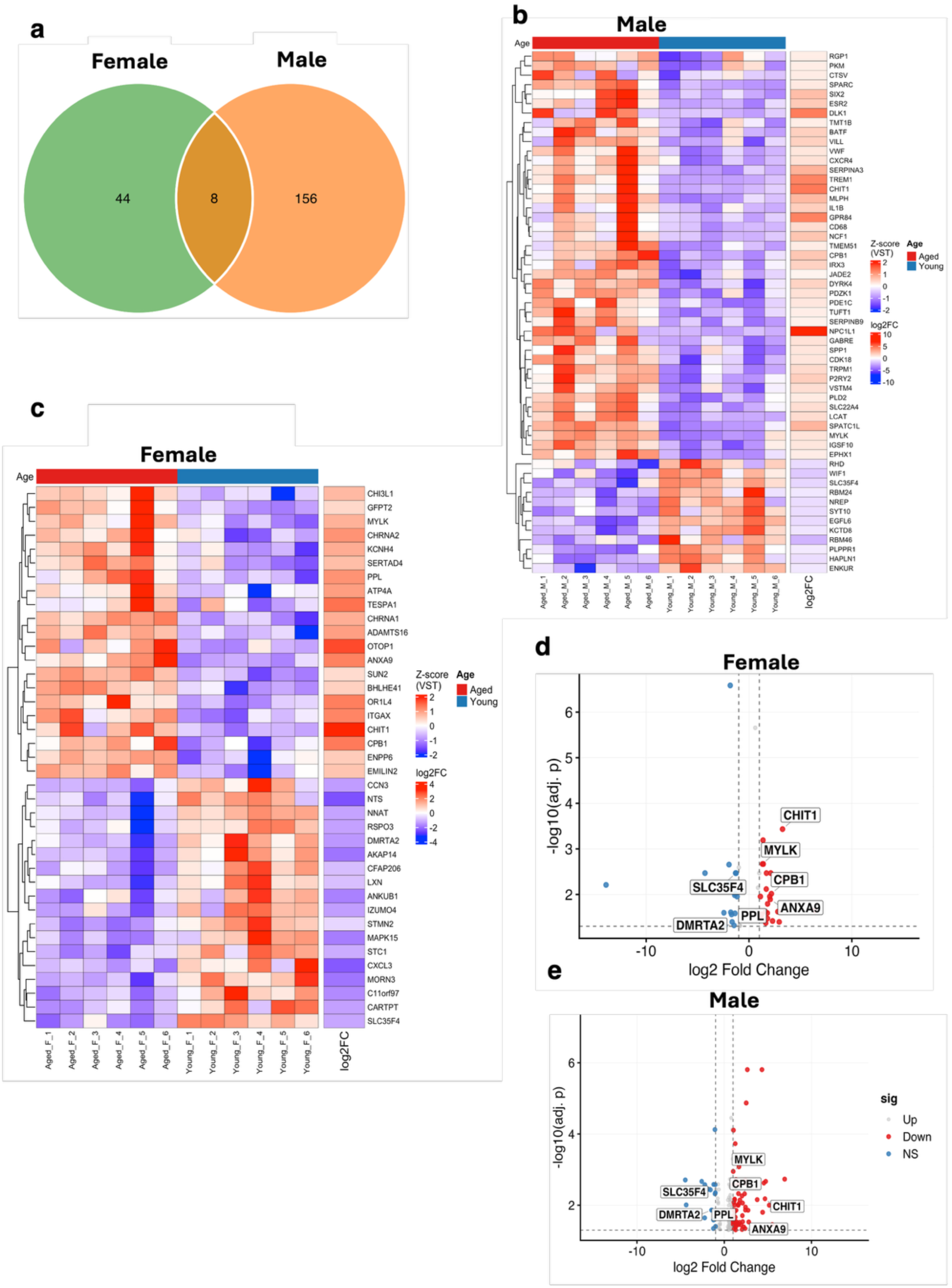
Age-Associated Differential Gene Expression Reveals Sex-Dependent Expression Patterns. **a**, Venn diagram highlighting convergence of DEGs between sex groups. **b**, Heatmap showing significantly differentially expressed genes (DEGs; adjusted *P* ≤ 0.05) identified from male samples. Genes are ranked by statistical significance and displayed as row-wise *z*-score–normalized expression values across individual samples. Columns represent samples ordered by age group, with age group and sex indicated by the top annotation bar. K-means clustering was applied to genes to highlight shared expression patterns, while samples were not clustered to preserve biological ordering. A side heatmap indicates the corresponding log₂ fold change (aged versus young), with red denoting increased expression and blue denoting decreased expression with age. **c,** Heatmap showing significantly differentially expressed genes (DEGs; adjusted *P* ≤ 0.05) identified from female samples, displayed and interpreted as in **b. d**, Volcano plot showing gene-level differential expression between aged and young males with the 8 DEGs shared between sexes denoted in **a** labeled. Each point represents a gene, plotted by log₂ fold change (aged versus young) and −log₁₀–transformed adjusted *P* value. Genes with adjusted *P* ≤ 0.05 and absolute log₂ fold change ≥ 1 are highlighted in red (upregulated) or blue (downregulated), while genes with log₂ fold change < 1 are shown in grey. Dashed lines indicate statistical significance and fold-change thresholds, and selected genes are labeled. **e**, Volcano plot showing gene-level differential expression between aged and young females with the 8 DEGs shared between sexes denoted in **a** labeled, displayed and interpreted as in **d.**

### Female aging is associated with glial reactivity and inflammatory remodeling

Several female-specific differentially expressed genes were associated with microglial activation, reactive glial states and inflammatory signaling, suggesting age-related remodeling of the female hippocampal glial microenvironment (Fig. 2c). Notably, this signature included *CHIT1*, *CHI3L1* (YKL-40) and *ITGAX* (CD11c), which have been associated with activated or reactive microglial states^43^, together with the chemokine *CXCL3* (Fig. 2c). CHIT1 has been implicated in microglial phagocytic and inflammatory responses, while ITGAX/CD11c is commonly used as a marker of activated microglial populations. Consistent with this pattern, *CHIT1* (+3.25, padj = 3.7 × 10⁻⁴), *CHI3L1* (+1.18, padj = 0.034), and *ITGAX* (+2.01, padj = 0.011) were increased with age. However, the response was not uniformly pro-inflammatory, as *CXCL3* was reduced (−2.34, padj = 0.048), together with *STC1* (−1.16, padj = 0.011) and *CCN3* (−1.39, padj = 0.010) (Supplementary Table 4). Thus, female hippocampal aging was characterized by selective remodeling of glial-associated transcriptional programs, with increased expression of genes associated with reactive and phagocytic microglial states alongside reduced expression of specific chemokine and tissue-remodeling-associated transcripts.

### Shared age-associated genes show largely concordant regulation across sexes

Eight genes were significantly age-associated in both females and males: *CHIT1, MYLK, PPL, ANXA9, CPB1, SLC35F4, DMRTA2,* and *LXN*. Most shared genes exhibited concordant directions of age-associated change between sexes (Fig. 2d,e), with several showing similar fold changes (Supplementary Table 5). *SLC35F4* decreased in both females and males (log₂FC = −1.25 and −1.22, respectively), whereas *ANXA9* (+2.06 and +2.23) and *CPB1* (+2.18 and +2.03) increased in both sexes. *DMRTA2* also showed nearly identical decreases in females and males (log₂FC = −1.45 in both). In contrast, *LXN* displayed sex-divergent regulation, decreasing in females (log₂FC = −0.89, padj = 0.048) but increasing in males (log₂FC = +0.67, padj = 0.035). Thus, although the overlap in age-associated genes between sexes was limited, the majority of shared genes showed concordant age-associated regulation, with *LXN* representing a notable exception.

### Shared and sex-specific inflammatory pathway enrichment across sexes

To assess whether specific biological pathways are associated with aging across sexes, we performed gene set enrichment analysis (GSEA) against the MSigDB Hallmark collection as per the literature^44^. Twenty Hallmark sets reached significance in at least one sex, and the majority were positively enriched in the aged condition (Fig. 3a). Aging in both sexes was associated with enrichment of hallmark inflammatory gene sets, including HALLMARK_ IL2_STAT5_SIGNALING, HALLMARK_INFLAMMATORY_RESPONSE, INTERFERON-γ_RESPONSE, and HALLMARK_COMPLEMENT(Fig. 3a), indicating a shared age-associated activation of immune programs. In both sexes, aging was additionally linked to enrichment of HALLMARK_P53_PATHWAY (Supplementary Fig. 2a-c), HALLMARK_INTERFERON_ALPHA_RESPONSE, HALLMARK_TNFA_SIGNALING_VIA_NFKB, and, HALLMARK_IL6_JAK_STAT3_SIGNALING (Fig. 3b-c).

**Fig 3.**
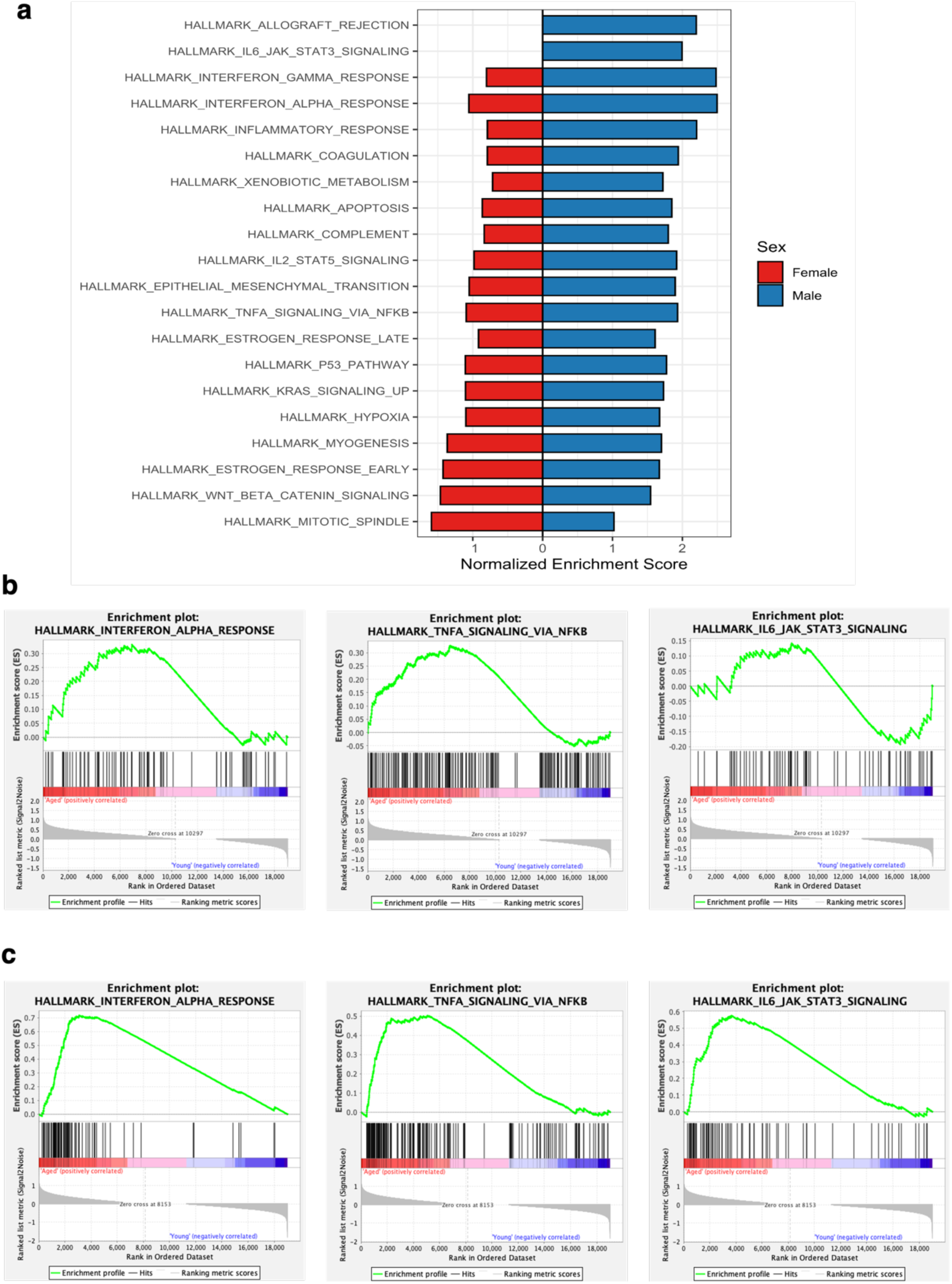
Aging Is Associated with Shared Inflammatory Pathway Enrichment and Sex-Specific Metabolic and Hormonal Signatures. **a,**Bar plot summarizing Hallmark gene set enrichment results from gene set enrichment analysis (GSEA) performed separately in females and males using default parameters in the GSEA application. Pathways shown represent the top enriched gene sets (top n = 20) with a false discovery rate (FDR) ≤ 0.25 in at least one sex, ranked by maximum absolute normalized enrichment score (NES). Bars indicate the NES for each pathway, with enrichment in males shown to the right (positive NES) and enrichment in females mirrored to the left for visual comparison. **b,** Representative GSEA enrichment plot for a top-ranked gene set in males comparing aged males (Old_M) versus young males (Young_M). The running enrichment score (green line) reflects the accumulation of pathway genes across the ranked gene list, ordered by differential expression with age. Vertical black bars indicate positions of gene set members within the ranked list. Positive enrichment indicates higher expression in aged males relative to young males. **c,** Representative GSEA enrichment plot for a top-ranked gene set in females comparing aged females (Old_F) versus young females (Young_F), displayed as in **b**.

### Estrogen-response enrichment is similar between sexes but driven by distinct genes

Estrogen signaling has been implicated in neuroprotection.^45^ Loss of estrogen in the brain has been shown to accelerate amyloid-β (Aβ) plaque deposition in Alzheimer’s disease,^46^ while estrogen replacement therapy reduced Aβ plaque accumulation in aged female rhesus macaques^47^. In females, brain estrogen levels decline significantly following menopause, a change that has been proposed to contribute to the increased susceptibility of women to developing Alzheimer’s disease^48, 49, 50^. Both sexes also exhibited enrichment of early and late estrogen response pathways (Fig. 3a)(Supplementary Fig. 4a-c), indicating increased engagement of estrogen-related signaling in the aging brain. The early estrogen response was enriched in aged animals of both sexes to a similar extent, with peak enrichment scores of 0.40 (Supplementary Fig. 4b-c). Leading-edge genes were distributed throughout the age-associated portions of the ranked gene lists, with the zero-crossing of the ranking metric occurring at rank 10,297 in females and 8,153 in males(Supplementary Fig. 4b-c). Thus, at the pathway level, estrogen-response enrichment appeared largely comparable between sexes. However, hierarchical clustering of the leading-edge genes across all 16 samples revealed substantial differences in the genes contributing to this enrichment (Supplementary Fig. 4a). In females, 40 genes comprised the leading-edge subset, including genes involved in lipid and sterol metabolism, cyclic-nucleotide signalling, transcriptional regulation, and neuronal function. Prominent contributors included *FASN, ELOVL2, ELOVL5, ABCA3,* and *SULT2B1*, together with the adenylate cyclases *ADCY1* and *ADCY9* and the scaffolding protein *AKAP1*. Transcriptional and chromatin regulators, including *NCOR2, RARA, KLF10, CBFA2T3*, *KDM4B, MED13L*, and *FOXC1,* as well as neuronal and cytoskeletal genes, including *MAPT, CELSR2, SYT12, SYNGR1, SYBU,* and *TIAM1*, also contributed to the female enrichment (Supplementary Fig. 4a).

In males, 35 genes contributed to the same hallmark but showed a distinct leading-edge composition. The male subset included *KRT19, CLDN7, SLC9A3R1, MYOF, TGM2, SCNN1A, ESRP2, ELF3, ELF1,* and *TFAP2C,* together with proliferation- and cellular stress-associated genes including *MYC, MYB, RBBP8,* and *XBP1*. Additional contributors included genes related to sterol metabolism and redox homeostasis, such as *DHCR7, NADSYN1, BLVRB, DHRS3,* and *PEX11A,* as well as *IGFBP4* and *CISH*, which participate in growth-factor and cytokine signalling. Thus, although both sexes showed enrichment of the same estrogen-response hallmark, the leading-edge genes contributing to this enrichment differed substantially, indicating that a shared pathway-level signature can be underpinned by distinct transcriptional programs during hippocampal aging.

### Sex-specific contributors to inflammatory pathway enrichment

To determine whether pathways enriched in both sexes were driven by shared or distinct transcriptional components, we examined the core-enrichment genes underlying each GSEA result (Fig. 4). For the interferon-α response, 29 genes contributed to enrichment in both sexes, including *ADAR, IFIT2, IFIT3, IL15, IRF9, MX1,* and *OASL*, defining a shared interferon-responsive core (Fig. 4a). However, 11 genes were female-specific and 35 were male-specific. Female-specific contributors included *IL4R, STAT2,* and *TRIM25*, whereas the male-specific subset included *CD74, CXCL10, ISG15, PSMB9, SP110,* and *USP18*, indicating distinct transcriptional contributions to the shared pathway-level response. A similar pattern was observed for TNFα/NF-κB signalling, with 51 core-enrichment genes shared between sexes, including *FOS, JUN, NFKB1, RELA, PTGS2,* and *TNFAIP3* (Fig. 4b). An additional 29 genes were female-specific and 61 male-specific. The female-specific subset included *EGR1, KLF10, NFE2L2,* and *SMAD3*, whereas the male-specific subset contained several inflammatory mediators and regulators, including *CCL2, IL1B, IL6, IL23A, TNF, TLR2,* and *SOCS3*. The IL6/JAK/STAT3 pathway showed the greatest divergence, with only seven core-enrichment genes shared between sexes, including *CXCL9, CXCL10, CXCL11, IL2RA,* and *IRF1* (Fig. 4c). Eight genes contributed exclusively in females, including *IFNGR1, IL6,* and *IL7*, compared with 37 male-specific genes, including *IL1B, IL1R1, MYD88, STAT1, STAT3, TNF,* and *TYK2*. Together, these analyses show that shared pathway-level enrichment does not reflect a uniform transcriptional response across sexes. Although aging converged on common interferon, TNFα/NF-κB, and IL6/JAK/STAT3 pathways, the genes contributing to these responses were substantially sex-specific, with particularly pronounced divergence in the IL6/JAK/STAT3 pathway.

**Fig. 4.**
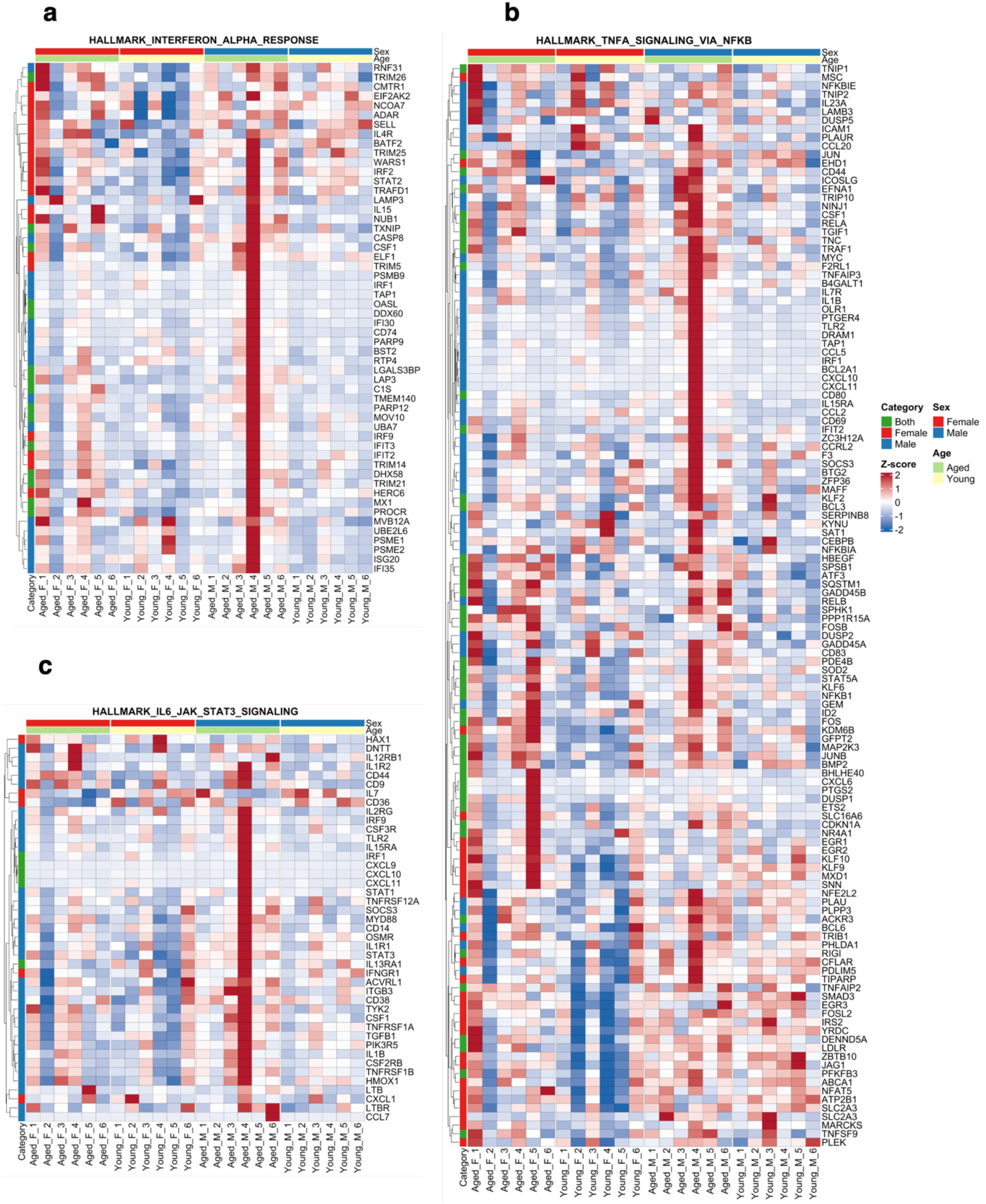
Sex-specific and shared gene-level contributors to age-associated inflammatory pathway enrichment. **a–c**, Heatmaps depicting gene-level contributions to age-associated enrichment of inflammatory Hallmark pathways identified by gene set enrichment analysis (GSEA). Panels show Hallmark interferon alpha response **(a)**, Hallmark interferon gamma response **(b),** and Hallmark complement **(c).** For each pathway, rows represent genes from the GSEA ranked list that contribute to the enrichment signal, restricted to core enrichment genes (CORE.ENRICHMENT = “Yes”). Genes are categorized based on whether they contribute to pathway enrichment in females only, males only, or both sexes, as indicated by the row-side annotation. Categories were defined by the overlap of core enrichment genes between sex-stratified GSEA results. Columns represent individual samples ordered by sex and age group, with sex and age indicated in the top annotation bar. Gene expression values are shown as row-wise z-score–normalized expression, enabling comparison of relative expression patterns across samples. Hierarchical clustering was applied to genes to highlight shared expression dynamics.

### Gene co-expression modules associated with age in females and males

To identify transcriptional networks associated with ageing, we performed weighted gene co-expression network analysis (WGCNA), identifying 17 co-expression modules (Supplementary Table 15). Correlation of module eigengenes with age–sex traits identified 14 modules significantly associated with ageing in a sex-specific manner (P < 0.05; Fig. 5b). In females, MEblue and MEbrown were positively associated with age, whereas MEturquoise, MElightcyan and MEmidnightblue were negatively associated. In males, MEgreenyellow, MEsalmon, MEgrey60, MEred, MEcyan and MEtan were positively associated with age, while MEmagenta, MEgreen and MEpink were negatively associated (Fig. 5b). Hierarchical clustering further revealed distinct network organization associated with female and male ageing, with the aged-female trait clustering with MEblue and MEbrown and the aged-male trait clustering with a broader group of modules, including MEred, MEsalmon, MEgreenyellow, MEcyan and MEtan (Fig. 5c). Pathway enrichment analysis identified distinct biological processes within the principal age-associated modules (Supplementary Fig. 5). In females, the age-increased MEblue module was enriched for cellular senescence and stress-related pathways, including oxidative stress- and oncogene-induced senescence (Supplementary Fig. 5a-b; Supplementary Table 16) MEbrown, also positively associated with age, was enriched for synaptic membrane and neuronal signaling processes and additionally contained hormone- and stress-responsive signatures, including estradiol-response, HDAC-target and NRF2-associated programs(Supplementary Fig. 5c-d; Supplementary Table 17). In contrast, the age-decreased MEturquoise module showed strong enrichment for oxidative phosphorylation and mitochondrial energy metabolism, consistent with an age-associated reduction in a mitochondrial transcriptional network (Supplementary Fig. 5e-f; Supplementary Table 18). Together, these modules define a female ageing architecture characterized by reduced mitochondrial metabolic programs alongside increased senescence and remodeling of synaptic and stress-responsive networks. In males, the two age-associated modules with significant pathway enrichment, MEgreenyellow and MEcyan, were predominantly related to oligodendrocyte and myelin biology (Supplementary Fig 6). MEgreenyellow was enriched for oligodendrocyte marker genes, neuron ensheathment, peripheral myelination and Rho GTPase signaling, together with cell-cycle and vesicle-associated processes (Supplementary Fig 6a,b). MEcyan showed a more focused enrichment for oligodendrocyte differentiation, myelin sheath organization, axon ensheathment and structural constituents of myelin (Suplementary Fig 6c,d). Both modules increased with age, indicating coordinated remodeling of oligodendrocyte and myelin-associated transcriptional networks in the ageing male hippocampus. The remaining age-associated male modules did not show significant pathway enrichment under the applied threshold and were therefore not assigned pathway-level functions.

**Fig 5.**
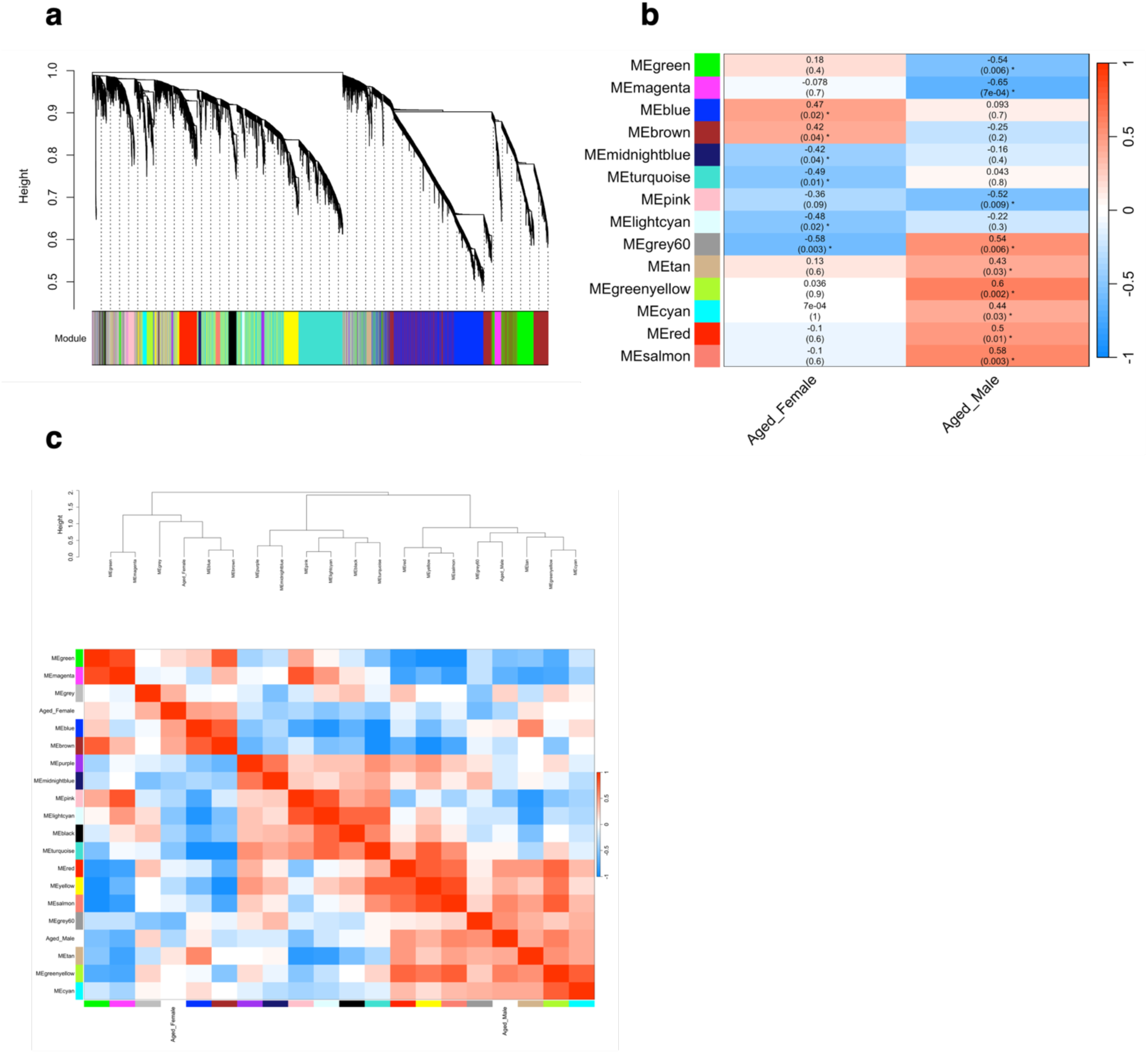
Gene co-expression modules associated with age- and sex-specific transcriptional states in rhesus. **a,**Hierarchical clustering of genes based on topological overlap identifies discrete co-expression modules, each labelled by a distinct colour. Modules were defined using dynamic tree cutting, and only high-quality samples and genes passing WGCNA quality control are shown. **b,** Heat map of Pearson correlations between module eigengenes and age–sex traits, highlighting modules significantly associated with aged females and aged males. The color scale denotes correlation strength and sign (red, positive; blue, negative), and each cell reports the correlation coefficient with the corresponding P value in parentheses; asterisks indicate significant associations (P < 0.05). **c,** Clustering of significant module eigengenes and traits with the corresponding eigengene adjacency heat map. The dendrogram summarizes eigengene similarity, and the adjacent heat map depicts pairwise eigengene correlations, where warm and cool colors indicate positively and negatively correlated eigengenes, respectively.

### Cross-species conservation of ageing-associated networks in Alzheimer’s disease

To determine whether the age-associated co-expression networks identified in rhesus hippocampus correspond to networks that change in the human context, we constructed a parallel WGCNA network from the human hippocampal single-cell transcriptomic dataset GSE268609^51^ that spanned young adults (Young Adult), healthy aged individuals (Healthy Aged), super agers (Super Agers), individuals with mild cognitive impairment (MCI), and individuals with Alzheimer’s disease (AD) and tested the resulting modules for overlap with the macaque modules. Clustering of human module eigengenes revealed distinct patterns of association with clinical traits (Supplementary Fig. 7a). Twenty modules were significantly associated with at least one trait, with the strongest associations observed for AD rather than normative ageing (Supplementary Fig. 7b). Eight modules were positively associated with AD, including MEgreen (r = 0.63, P = 1.7 × 10^5^), MEsaddlebrown (r = 0.62, P = 3 × 10^5^) and MEbrown (r = 0.59, P = 6.6 × 10^5^), with additional associations observed for MEmagenta, MEwhite, MEgrey60, MEdarkturquoise and MEcyan. Six modules were negatively associated with AD, including MEroyalblue (r = −0.59, P = 7.9 × 10^5^), MEblack (r = −0.56, P = 1.9 × 10^5^), MEturquoise (r = −0.50, P = 0.0011), MEorange (r = −0.48, P = 0.0018), MEblue (r = −0.46, P = 0.0034) and MEpink (r = −0.45, P = 0.004). In contrast, relatively few modules were associated with intermediate or normative ageing phenotypes. The young-adult trait was positively associated with MEdarkorange (r = 0.52, P = 7.5 × 10^4^) and MEdarkred (r = 0.33, P = 0.04), and negatively associated with MEgreen (r = −0.54, P = 4.2 × 10^4^), MEmagenta (r = −0.47, P = 0.0029), MEpaleturquoise (r = −0.46, P = 0.0033) and MEsaddlebrown (r = −0.38, P = 0.016). Only one module was associated with healthy ageing (MEdarkmagenta; r = 0.36, P = 0.026), and one with super-ager status (MElightyellow; r = 0.33, P = 0.039)(Supplementary Fig 7b; Supplementary Table 19). Notably, MEgreen, MEmagenta, and MEsaddlebrown showed opposing associations with young-adult status and AD, with lower eigengene values in young adults and higher values in AD. Together, these patterns indicate that, within this dataset, the biggest module-level transcriptional differences were associated with AD, whereas relatively fewer modules distinguished normative ageing or intermediate phenotypes.

### Age-associated macaque modules show similar gene membership in the human hippocampus

Because the rhesus network was structured around sex-specific ageing, whereas the human network was structured primarily around clinical phenotype, we assessed cross-species conservation based on gene membership rather than direct comparison of module–trait correlations. Gene overlap was quantified for every macaque–human module pair and evaluated using hypergeometric tests against the union of module genes (Supplementary Figs. 9 and 10; Supplementary Table 20). The cross-species overlap was extensive and non-random, with several macaque modules showing significant correspondence to human modules. The strongest intersection was observed between the macaque MEturquoise and human MEblue modules, comprising 553 shared genes (P < 0.001). Three additional intersections contained more than 200 genes, including macaque MEblue–human MEbrown (273 genes), macaque MEbrown–human MEbrown (205 genes), and macaque MEturquoise–human MEbrown (131 genes). The human MEbrown module also overlapped significantly with macaque MEgreenyellow (109 genes, P < 0.001), MEcyan (87 genes, P < 0.001), and MEsalmon (34 genes, P < 0.05). Additional significant correspondences included macaque MEred with human MEorange (81 genes, P < 0.001) and MEpaleturquoise (24 genes, P < 0.05); macaque MEbrown with human MEdarkturquoise (86 genes), MEroyalblue (30 genes), MEmagenta (25 genes) and MEturquoise (46 genes); macaque MEgreen with human MEdarkturquoise (41 genes, P < 0.001) and MEroyalblue (18 genes, P < 0.05); macaque MEpink with human MEblue (56 genes, P < 0.001); macaque MEsalmon with human MEorange (20 genes, P < 0.001); macaque MEtan with human MEdarkmagenta (7 genes, P < 0.001); and macaque MElightcyan with human blue (19 genes, P < 0.05) (Supplementary Fig. 10; Supplementary Table 20). Importantly, several of the macaque modules showing sex-specific associations with ageing corresponded to human modules altered in AD (Supplementary Fig 8). The macaque MEturquoise module, which was negatively associated with age in females, showed its strongest cross-species correspondence with the human MEblue module, which was negatively associated with AD (r = −0.46). Conversely, the macaque MEgreenyellow and MEcyan modules, both positively associated with age in males, showed significant overlap with the human MEbrown module, which was positively associated with AD (r = 0.59)(Supplementary Fig 8). These cross-species correspondences link sex-specific ageing-associated co-expression programs in the rhesus hippocampus with human hippocampal networks altered in AD, suggesting that distinct ageing trajectories may converge on transcriptional programs relevant to neurodegenerative disease.

### Conserved intersections are enriched for oxidative phosphorylation and myelination

We next tested the significantly overlapping gene sets for functional enrichment using the union of module genes as the background (Supplementary Fig. 11). The 553 genes shared between the macaque MEturquoise and human MEblue modules were enriched for five pathways, with HALLMARK_OXIDATIVE_PHOSPHORYLATION and KEGG_OXIDATIVE_PHOSPHORYLATION showing the strongest enrichment (60 and 44 genes, respectively; Supplementary Fig. 11a; Supplementary Table 21). The corresponding gene-concept network was characterized by a mitochondrial oxidative-phosphorylation program, with prominent representation of ATP synthase and cytochrome c oxidase components, including *ATP5F1D, ATP5F1E, ATP5MC1, ATP5ME, ATP5MF, ATP5MG, ATP5PB, ATP5PD, ATP5PO, COX4I1, COX5A, COX6A1, COX6B1, COX6C* and *COX8A* (Supplementary Fig. 11b). The remaining enriched terms, including REACTOME_REGULATION_OF_APOPTOSIS, REACTOME_TNFR2_NON_CANONICAL_NF_KB_PATHWAY and REACTOME_APOPTOSIS, were largely driven by a shared proteasome component together with apoptosis-related genes including *BAX, BAD, BCL2L1,* and *UBB* (Supplementary Fig. 11a-b). Thus, the strongest cross-species intersection represents a conserved mitochondrial and proteostatic program. Notably, this gene set links the macaque MEturquoise module, which was negatively associated with age in females, with the human MEblue module, which was negatively associated with AD (Supplementary Fig. 8).

The 109 genes shared between macaque MEgreenyellow and human MEbrown were enriched for ten pathways, with oligodendrocyte identity and differentiation among the dominant annotations, including LEIN_OLIGODENDROCYTE_MARKERS and GOBP_OLIGODENDROCYTE_DIFFERENTIATION (Supplementary Fig. 11c; Supplementary Table 22). The gene set also showed enrichment for BLALOCK_ALZHEIMERS_DISEASE_UP, providing an independent connection to an AD-associated transcriptional signature. The corresponding gene-concept network was characterized by a set of genes associated with oligodendrocyte differentiation and myelin biology, including *SOX10, CRYAB, FA2H, GJB1, GJC2, DNM2, PMP22, LITAF, MYH14, PLLP, TF, PACS2, LMNA, JUP,* and *CYP27A1* (Supplementary Fig. 11d). Similarly, the 87 genes shared between macaque MEcyan and human MEbrown were enriched for pathways related to oligodendrocyte differentiation, myelin structure and assembly, neuronal ensheathment and intracellular sterol transport (Supplementary Fig. 11e; Supplementary Table 23). These terms converged on a shared myelin- and oligodendrocyte-associated gene program, including *PLP1, MBP, MOBP, MOG, MAL, CNP, CLDN11, UGT8, ERMN, MYRF, PLLP, ASPA, CNTN2, GPR37, ABCA2, NPC1,* and *CYP27A1* (Supplementary Fig. 11f). Together, these analyses identify two major functional programs that exist in both macaque ageing-associated and human AD-associated networks. The female-associated macaque MEturquoise and human MEblue intersection was characterized predominantly by oxidative phosphorylation and proteostasis, whereas the male-associated macaque MEgreenyellow and MEcyan modules showed conserved oligodendrocyte- and myelin-associated programs through their overlap with the human MEbrown module. These findings connect sex-specific molecular signatures of hippocampal ageing in rhesus macaques with gene programs represented in the human AD transcriptome.

### Ovarian steroid-responsive genes partition into modules with opposing sex-specific age associations

Ovarian steroids regulate the expression of genes supporting serotonin and glutamate neurotransmission, synapse assembly, DNA repair, protein chaperoning, and axonal transport in the aged female macaque brain ^52^. To assess the network context of these hormone-responsive genes, we mapped each gene to its parent WGCNA module and examined the age associations of these modules in females and males (Supplementary Fig. 12).

The curated hormone-responsive gene panel mapped to five modules spanning distinct regions of the female–male age-correlation space. Eight genes mapped to MEturquoise, which was negatively associated with age in females (r = −0.49) but showed little association with age in males (r = 0.04). These included the chaperones *HSPD1, HSPB1*, and *HSP90B1*, the DNA-repair and genome-maintenance genes *NTHL1, PCNA,* and *RAD23A*, the transport-associated gene *DYNLL1*, and the neuronal protein *SNCA*. Four genes mapped to MEblue, which was positively associated with age in females (r = 0.47) but showed little association with age in males (r = 0.09), including the glutamate receptor *GRM1*, the neurotrophin receptor *NTRK2*, the corticotropin-releasing hormone receptor *CRHR1,* and the microtubule-associated protein *MAPT*.

Six genes mapped to modules showing opposing associations with age between sexes. In MEbrown, *GRIN2A, NCAM1,* and *ADAM10* mapped to a module positively associated with age in females (r = 0.42) but negatively associated with age in males (r = −0.25). Similarly, *SNAP25*, *GRIA2*, and *UBE3A* mapped to MEgreen, which showed a weak positive association with age in females (r = 0.18) and a stronger negative association in males (r = −0.54). Only *CRHR2* mapped to a module negatively associated with age in both sexes, MEpink (female r = −0.36; male r = −0.52). No genes in the curated panel mapped to a module positively associated with age in males and negatively associated with age in females.

The distribution of these genes across modules revealed distinct functional and age-associated network contexts. Chaperone and DNA-repair genes were concentrated within MEturquoise, whereas glutamate receptor- and synapse-associated genes, including *GRM1, GRIN2A, GRIA2, NTRK2, NCAM1,* and *SNAP25*, mapped to modules positively associated with age in females. Overall, ovarian steroid-responsive genes were distributed across multiple co-expression programs with distinct relationships with age between sexes. This distribution highlights the sex-specific organization of hormone-responsive genes within the ageing hippocampal transcriptome.

### Sex-specific isoform remodeling with aging

To resolve age-related change at the level of individual transcripts rather than summarized gene expression, we tested for differentially expressed isoforms (DIEs) between aged and young animals within each sex (Fig. 6). The two sexes yielded markedly different numbers of significant isoforms, and the sets were almost entirely disjoint. We identified 526 DIEs in females and 21 in males, of which only two were shared (Supplementary Table 8): an isoform of *SERPINA3* (ENSMMUT00000039534) and an isoform of *APOC2* (ENSMMUT00000106127) (Fig. 6a). Both were increased with age in each sex (Fig. 6d, e), making them the only transcripts in this analysis with a concordant age response in males and females. Isoform-level testing resolved several genes into multiple independently significant transcripts. Nine isoforms of the astrocytic chloride channel *TTYH1* were decreased in aged females (log₂ fold change −1.57 to −1.73; adjusted P = 1.5 × 10⁻⁵ to 0.11), and multiple isoforms were likewise recovered for *SERPINA1* (four isoforms, log₂ fold change +1.59 to +1.70), *CCDC13* (five isoforms, +1.12 to +2.00), *COMT* (eight isoforms, +0.75 to +0.95), *FEZ1* (four isoforms, +0.77 to +0.87), *DENND2A* (six isoforms, −1.78 to −2.17) and *ITM2A* (three isoforms, +1.22)(Supplementary table 6).

**Fig 6.**
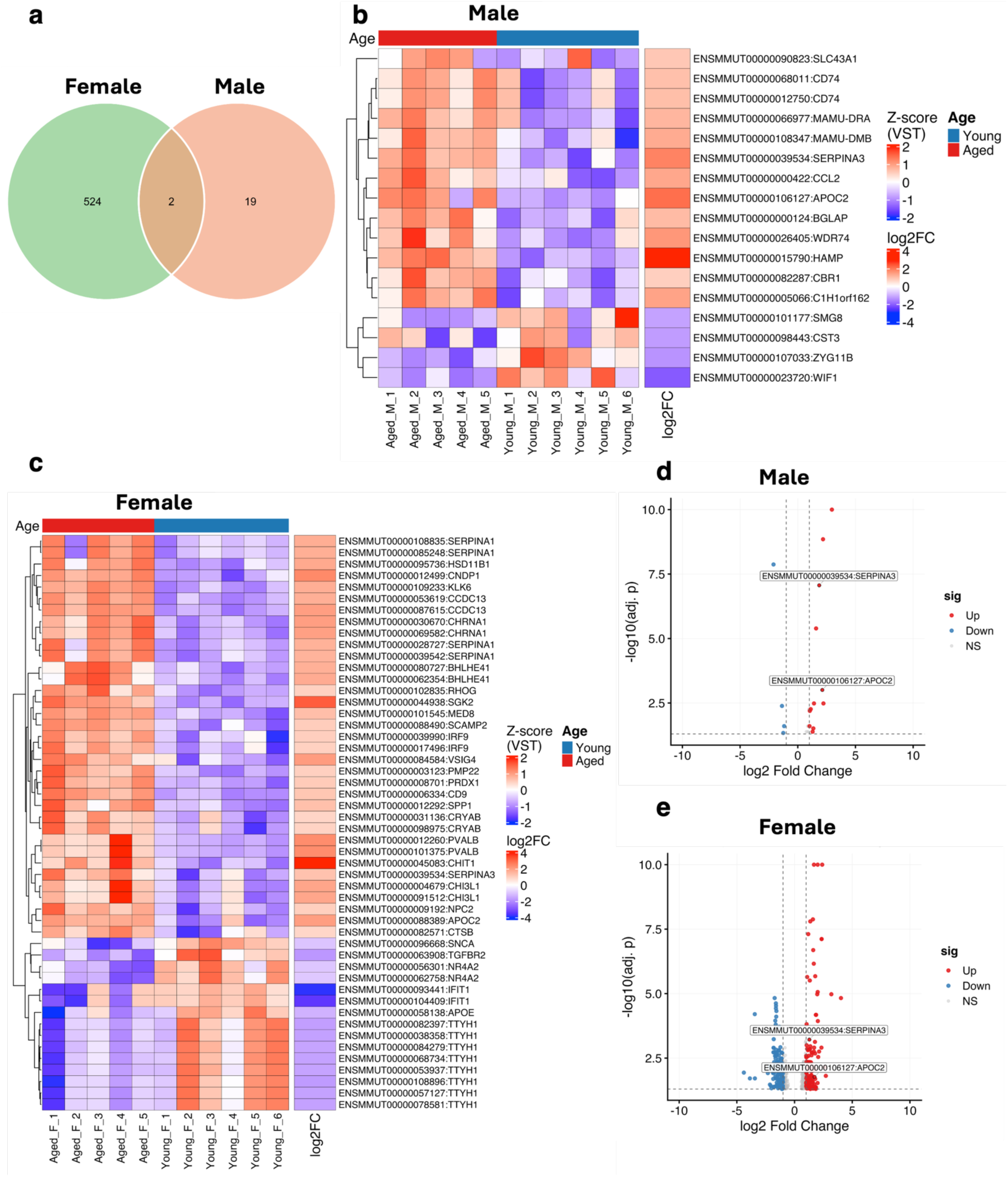
Sex-specific isoform usage reveals age-dependent transcriptional divergence between female and male rhesus macaques. **a,**Venn diagram highlighting convergence of differentially expressed isoforms (DIEs) between sex groups**. b,** Heat map of Z-score–normalized expression values for the top differentially expressed isoforms only in males, selected by adjusted *P* value with additional isoforms of interest included where indicated. Rows represent individual isoforms (labelled by gene name and feature identifier), columns represent male samples relabelled with anonymized IDs, and samples are ordered by sex and age metadata. A top annotation bar indicates age (Young, blue; Aged, red) and the right-hand side bar encodes isoform log2 fold change between the contrasted conditions (blue, downregulated; white, unchanged; red, upregulated). c, Heat map of Z-score–normalized expression values for the top differentially expressed isoforms in females, selected by adjusted *P* value with additional isoforms of interest included where indicated. Rows represent individual isoforms (labelled by gene name and feature identifier), columns represent all samples relabelled with anonymized IDs, and samples are ordered by age group. A top annotation bar indicates age (Young, blue; Aged, red), and the right-hand side bar encodes isoform log2 fold change between the contrasted conditions (blue, downregulated; white, unchanged; red, upregulated). **d**, Volcano plot showing isoform-level differential expression between aged and young males with the 2 DIEs shared between sexes denoted in **a** labeled. Each point represents an isoform, plotted by log₂ fold change (aged versus young) and −log₁₀–transformed adjusted *P* value. Isoforms with adjusted *P* ≤ 0.05 and absolute log₂ fold change ≥ 1 are highlighted in red (upregulated) or blue (downregulated), while isoforms with log₂ fold change < 1 are shown in grey. Dashed lines indicate statistical significance and fold-change thresholds, and selected isoforms are labeled. **e**, Volcano plot showing isoform-level differential expression between aged and young females with the 2 DIEs shared between sexes denoted in **a** labeled, displayed and interpreted as in **d.**

In females, the most significant DIEs were increased with (Fig. 6c). The strongest was *CNDP1* (log₂ fold change +2.39; adjusted P = 3.5 × 10⁻¹⁵), followed by *KLK6* (+1.68; P = 3.0 × 10⁻¹³), *CCDC13* (+2.00; P = 3.2 × 10⁻¹²), *SERPINA1* (+1.59; P = 1.3 × 10⁻⁸), *MED8* (+1.36), CD9 (+1.19) and *PVALB* (+2.34)(Fig 6c; Supplementary table 6). The largest fold changes belonged to the macrophage chitinase *CHIT1* (+4.02) and *SGK2* (+3.19). Also increased were the myelin and oligodendrocyte transcripts *MBP, MOG, PMP22, PMP2, CMTM5, QDPR,* and *DBNDD2*, the chaperone *CRYAB*, and a group of inflammation-associated and lysosomal genes including *CHI3L1, SERPINA3, SPP1, NPC2, CTSB, GM2A, FTL, VSIG4,* and the MHC class II transcript *MAMU-DRB1*(Supplementary table 6). Decreased isoforms were enriched for astrocytic and vascular transcripts *TTYH1, SLC1A3, GJA1, CLDN5, EGFL7, VWF, SPARC* and *ADGRG1* together with *APOE* (−1.33), *SNCA* (−1.02), *GRIA2, NRXN1, CALM3* and *CTNNB1* (Supplementary table 6; Fig 6c). The female signature therefore describes a coordinated shift toward glial activation, myelin-associated and lysosomal transcripts, with reduced astrocytic, synaptic and endothelial representation. Twenty-one isoforms were significant in males, of which 19 were male-specific (Fig. 6b). The set was led by the iron-regulatory peptide *HAMP* (log₂ fold change +2.96; adjusted P = 3.5 × 10⁻¹⁷), by a wide margin the most significant isoform in either sex, followed by an unannotated transcript (ENSMMUG00000029122, +2.19), *WIF1* (−2.11; P = 1.3 × 10⁻⁸), *SERPINA3* (+1.86) and *WDR74* (+1.59). Increased isoforms were dominated by antigen presentation and secreted immune factors: two isoforms of the invariant chain *CD74* (+0.97 and +0.99), the MHC class II transcripts *MAMU-DRA* (+1.12) and *MAMU-DMB* (+1.28), the chemokine *CCL2* (+1.35), *APOC2* (+2.13), *C1H1orf162* (+1.40), *BGLAP* (+1.04), *SLC43A1* (+0.86), and *CBR1* (+0.74) (Supplementary table 7; Fig 6b). Four isoforms were decreased: the Wnt antagonist *WIF1, ZYG11B* (−1.37), *SMG8* (−1.18), and the cystatin *CST3* (−1.26), and three of the 21 transcripts mapped to unannotated loci (ENSMMUG00000029122, represented by two isoforms, ENSMMUG00000031232 and ENSMMUG00000030326) (Supplementary table 7; Fig 6b). Antigen presentation and secreted immune and iron-handling factors, therefore, account for most of the male age response detectable at isoform resolution.

Much of this signal was not recoverable from conventional gene-level analysis. Plotting each of the top 40 isoforms alongside the expression of its parent gene in the same animals (Fig. 7) shows the age-related contrast attenuated or abolished once transcripts from the same gene are aggregated: the isoform triangles separate aged from young animals for essentially every row, while the corresponding gene triangles frequently do not, and the gene-level fold changes are compressed toward zero relative to the isoform-level values for the same transcripts. Gene-level significance was confined to a minority of them: *CHIT1, SGK2,* and *TESPA1*, and the *IFIT1* transcripts in females (Fig. 7a) and *APOC2* and *SERPINA3* in males (Fig. 7b). The remainder, including *CNDP1, PVALB, CHI3L1,* and *DENND2A* in females, and *HAMP, CD74,* and the MHC class II transcripts in males, were significant only at the isoform level. *HAMP,* the most significant isoform in either sex, was among those not recovered as a significant gene. Aggregating transcripts to the gene, therefore, discards a substantial fraction of the age-related signal in this dataset, and much of the sex-specific response described above is invisible to gene-level testing. Together, these findings demonstrate that hippocampal aging is characterized by sex-dependent remodeling of transcript isoform expression, with females exhibiting a substantially larger repertoire of age-associated isoform changes than males. This highlights the importance of transcript-level analyses for resolving molecular mechanisms of brain aging that are not evident from gene-level expression alone.

**Fig 7.**
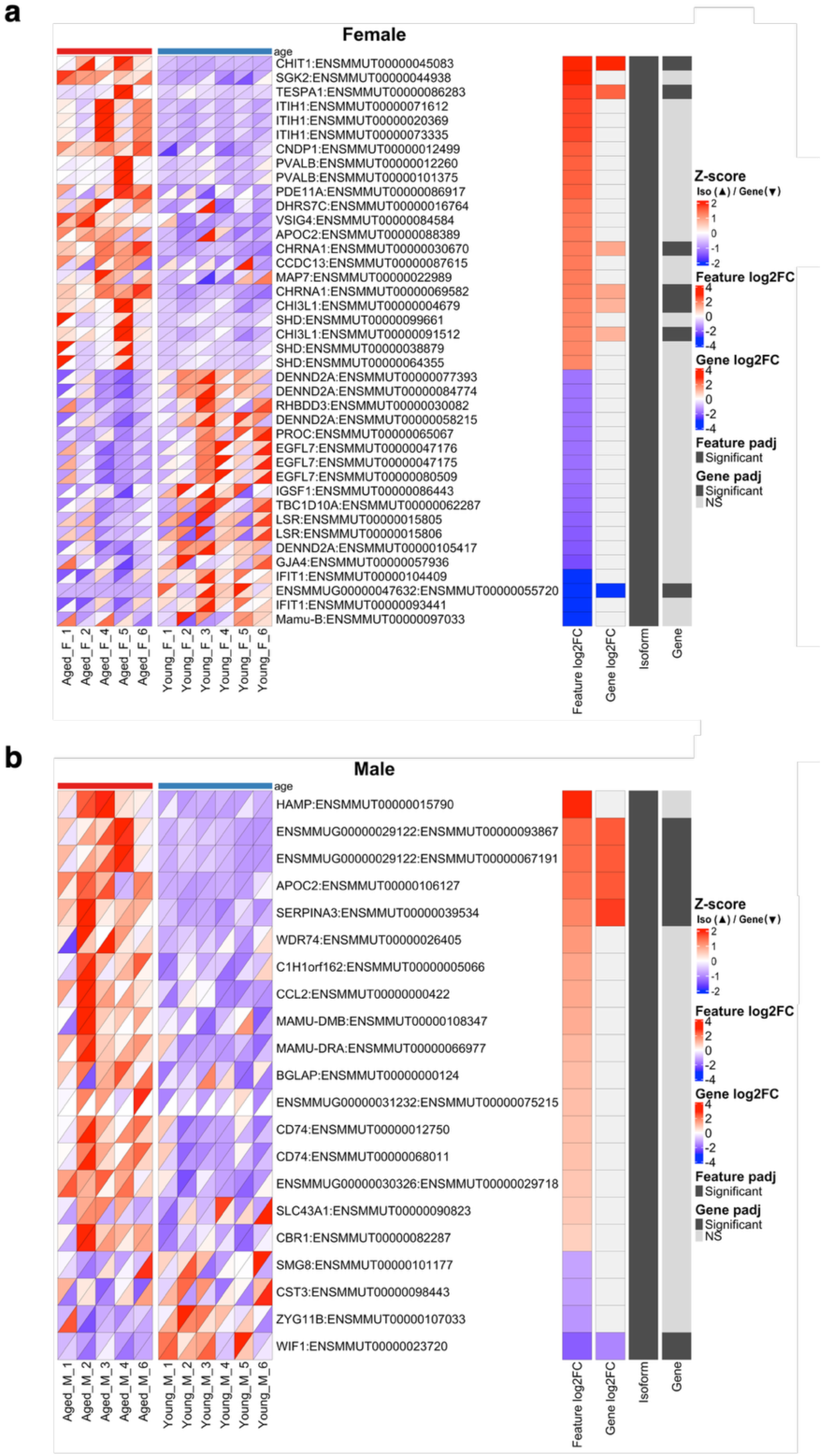
Isoform-specific expression changes reveal sex differences not detected at the gene level. **a,**Combined isoform and gene expression heatmap for females, showing the top 40 differentially expressed isoforms (rows) across samples (columns). For each feature, the upper triangle of each cell represents z-score–normalized isoform expression, and the lower triangle represents the corresponding gene-level expression. Rows are ordered by isoform-level differential expression. Adjacent annotations indicate isoform and gene log₂ fold change and adjusted p-values. **b,** Combined isoform and gene expression heatmap for males described and interpreted as in **a.**

### Differential isoform usage with age across sex

Isoform ratios are a determinant of cellular proteoform composition, as the relative abundance of distinct transcripts may dictate which protein species predominates within a given cellular state. Alternative splicing of a single gene can generate multiple isoforms and resulting proteoforms that differ in key functional domains. Shifts in isoform usage can therefore modulate protein localization, interaction networks, and aggregation propensity. The protein product encoded by the most abundant or functionally dominant transcript is commonly referred to as the canonical isoform; disruption of the balance between canonical and non-canonical isoforms can reprogram downstream signaling pathways and cellular stress responses. To determine whether aging alters isoform usage in a sex-specific manner, we performed differential transcript usage (DTU) analysis with IsoformSwitchAnalyzeR on isoform-level counts, testing aged against young animals within each sex. 30 genes showed significant age-associated isoform switching in females and 3 in males (gene-level switch q < 0.05)(Supplementary tables 9-10), contributing 81 and 12 switching isoforms, respectively (Supplementary tables 11-12) and paralleling the asymmetry seen in isoform-level differential expression (DIE). We first classified the event consequences of each switch. In both sexes, the switches arose predominantly from alternative transcript start and end sites. In females, alternative transcription end sites (41 isoforms) and alternative transcription start sites (40 isoforms) were the two most frequent consequences, followed by exon skipping (20), multiple exon skipping (18), alternative 3′ splice sites (7), intron retention (6), alternative 5′ splice sites (5) and mutually exclusive exons (2) (Fig. 8a).

**Fig 8.**
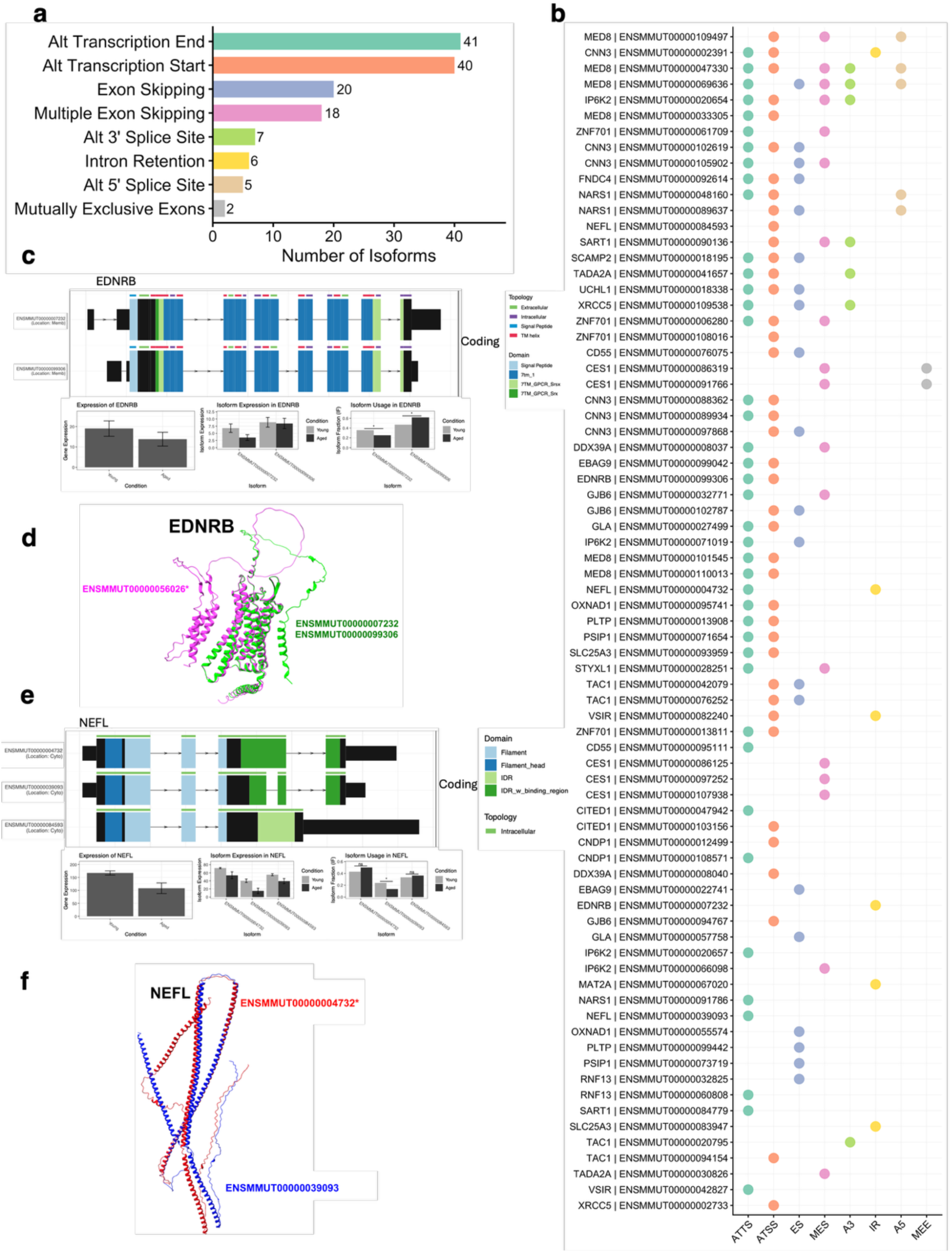
Isoform-level analysis reveals age-associated transcript usage changes in the female rhesus macaque hippocampus. **a,**Number of isoforms associated with each class of alternative splicing event detected between young and aged female hippocampi. **b,** Isoforms showing significant age-associated changes in usage. Each point represents an event detected for an isoform, colored by event class: alternative transcription termination site (ATTS), alternative transcription start site (ATSS), exon skipping (ES), multiple exon skipping (MES), alternative 3′ splice site (A3), intron retention (IR), alternative 5′ splice site (A5) and mutually exclusive exons (MEE). Isoforms are labeled by gene symbol; individual isoforms may contain multiple event classes. **c,** Representative isoform switch in endothelin receptor type B (EDNRB). Transcript models show exon structure, annotated Pfam domains, predicted topology and coding status for isoforms preferentially used in young and aged animals. Bar plots show gene-level expression, isoform-level expression and isoform usage (isoform fraction, IF) by age group. **d,** AlphaFold-predicted structures of the two EDNRB proteoforms superimposed, showing the canonical isoform ENSMMUT00000056026 in magenta and non canonical isoforms ENSMMUT00000007232 and ENSMMUT00000099306 in green. **e,** Representative isoform switch in neurofilament light chain (NEFL), presented as in **c.** Transcript models show filament and filament-head domains, intrinsically disordered regions (IDRs) and predicted binding regions. **f,** AlphaFold-predicted structures of the two NEFL proteoforms superimposed, showing the canonical isoform ENSMMUT00000004732 in red and non canonical ENSMMUT00000039093 in blue. Isoform switches were identified using IsoformSwitchAnalyzeR (young, n = 6; aged, n = 5). In **c** and **e,** bars show mean values and error bars indicate 95% confidence intervals. Statistical significance of isoform usage differences is indicated by *q < 0.05.

Resolving these consequences by isoform showed that individual transcripts frequently carry more than one splicing event (Fig. 8b). Forty-five of the 81 female isoforms and 7 of the 12 male isoforms were assigned two or more event types, and one female isoform carried five, indicating that switches commonly involve coordinated changes at both ends of the transcript as well as internal exon usage (Fig 8b; Supplementary tables 11-12). In females, the switching genes were led by *MED8* (q = 4.2 × 10⁻¹⁹) and *RNF13* (q = 3.4 × 10⁻¹³) and included *CITED1, SLC25A3, UCHL1, SCAMP2, VSIR, OXNAD1, PSIP1*, *CES1*, *EDNRB*, and *NEFL*, several of them represented by multiple switching isoforms (Fig 8b). Isoform switches illustrate a regulatory change that gene-level quantification cannot capture (Fig. 8c-f). In females, EDNRB (q = 0.0033), usage shifted between two coding isoforms, with ENSMMUT00000007232 decreasing and ENSMMUT00000099306 increasing in isoform fraction with age, against only a modest reduction in total gene expression (Fig 8c). Both isoforms are predicted to localize to the membrane and share the domain architecture of a class A G protein-coupled receptor: an N-terminal signal peptide, the 7tm_1 domain, and seven predicted transmembrane helices. Their coding sequences are largely the same, and they differ mainly in their terminal exons. ENSMMUT00000099306 uses alternative transcription start and end sites, and ENSMMUT00000007232 carries an intron-retention event and an extended 3′ terminal exon (Fig 8b,c). With age, ENSMMUT00000007232 expression fell by roughly half while ENSMMUT00000099306 stayed nearly stable. As a result, ENSMMUT00000007232 usage decreased (IF ≈ 0.37 to 0.25) and ENSMMUT00000099306 usage increased (IF ≈ 0.47 to 0.62). To visualize the predicted structures of these isoforms, we generated AlphaFold models and compared them with the reference isoform ENSMMUT00000056026 (Fig 8d).

ENSMMUT00000007232 and ENSMMUT00000099306 produced overlapping models, consistent with their shared coding sequence. Both resembled the reference model in the central seven-helix transmembrane bundle, with visible differences in the peripheral loop and helical regions. NEFL also showed reduced gene-level expression with age alongside isoform-specific changes (q = 0.034) (Fig 8e). Three coding isoforms were detected, all predicted to be intracellular. All three contain the N-terminal head domain and the central filament rod domain, and they differ at the C-terminus. ENSMMUT00000004732 encodes the full intrinsically disordered tail region with annotated binding motifs. In ENSMMUT00000039093, this region is interrupted and shortened. ENSMMUT00000084593 has an alternative 3′ structure that encodes a disordered segment without the annotated binding region. Total NEFL expression declined from 170 to 110 with age. ENSMMUT00000039093 showed the largest reduction and was the only isoform with a significant decrease in usage (IF 0.23 to 0.13). Usage of ENSMMUT00000004732 and ENSMMUT00000084593 did not change significantly.

AlphaFold models of ENSMMUT00000004732 (canonical) and ENSMMUT00000039093 are shown in Fig 8f. Both models show long α-helical segments corresponding to the rod domain, flanked by disordered terminal regions. The two isoforms differ mainly in the C-terminal region, reflecting the shortened tail of ENSMMUT00000039093. In MAMU-DMB, the most significant switch in males(q = 1.0 × 10⁻⁹), total gene expression was indistinguishable between young and aged animals, yet isoform usage was substantially reorganized. ENSMMUT00000040899 fell from approximately 0.63 to 0.36 in isoform fraction, and ENSMMUT00000108347 rose from approximately 0.20 to 0.40 (both P < 0.001), alongside a smaller significant increase in ENSMMUT00000040897 (Fig 9c). All three isoforms are predicted to localize to the membrane and contain a signal peptide and a C1-set domain. ENSMMUT00000040899 carries an MHC_II_beta domain, whereas ENSMMUT00000040897 and ENSMMUT00000108347 carry an MHC_II_alpha domain and differ from each other in their transcription start and end sites (Fig 9b,c). AlphaFold models of the three isoforms are shown in Fig 9d. All three share a β-sheet-rich core typical of Ig-like and MHC domains, with differences in the surrounding loops and terminal segments. In male BGLAP, where six isoforms were resolved, the age-related increase in gene expression was attributable to a single transcript (q = 1.2 × 10⁻⁹), ENSMMUT00000000124, whose usage rose from approximately 0.55 to 0.75 (P < 0.001) while the remaining five showed no significant change (Fig 9e). ENSMMUT00000000124 is predicted to be extracellular and encodes a signal peptide and the BGLAP_C domain, corresponding to secreted osteocalcin. The other coding isoforms instead contain mitochondrial carrier and/or Nnf1 domains and are predicted to localize to the mitochondria, cytoplasm, or nucleus.

**Fig 9.**
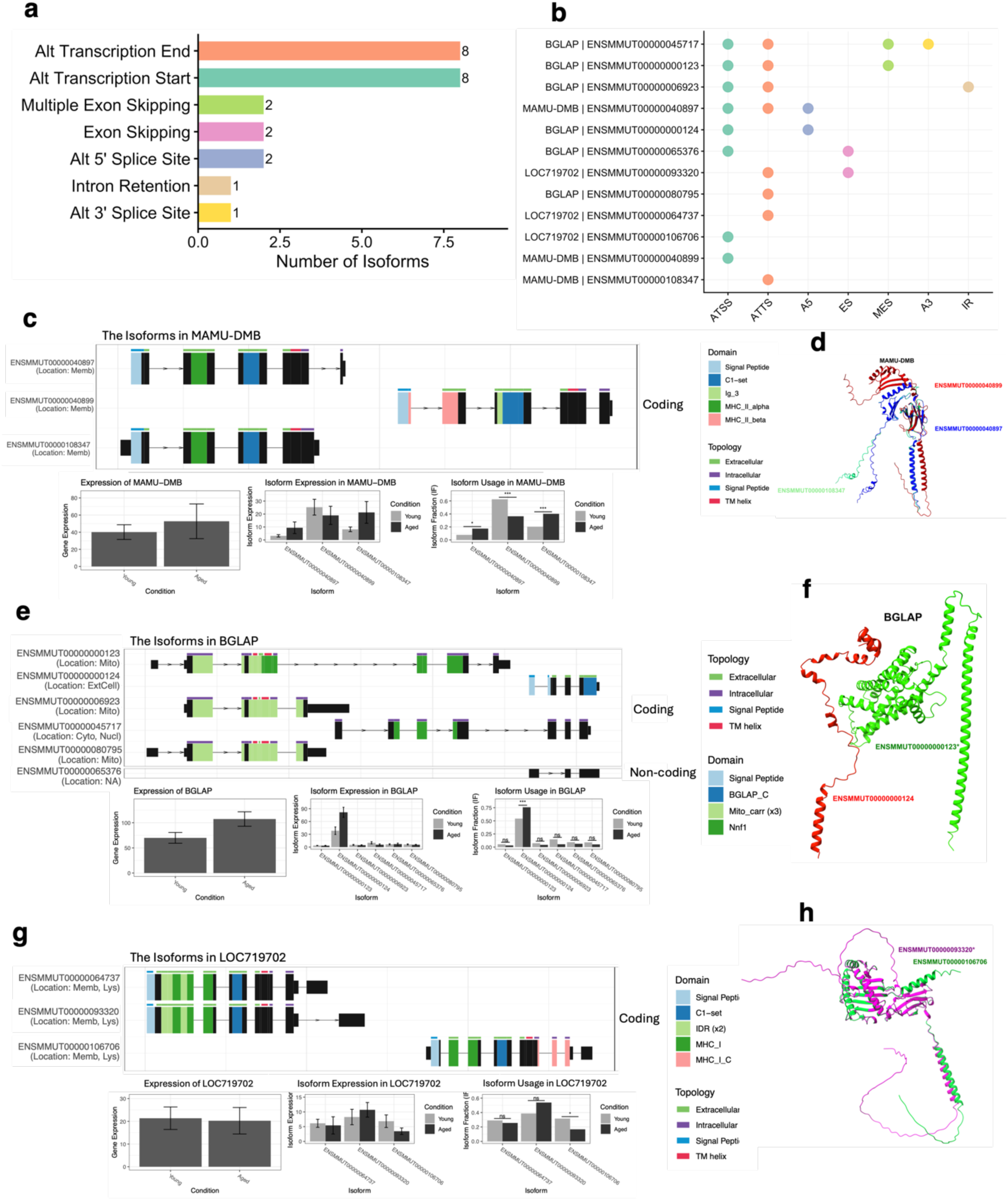
Isoform-level analysis reveals age-associated transcript usage changes in the male rhesus hippocampus. **a,**Number of isoforms associated with each class of alternative splicing event detected between young and aged male hippocampi, including alternative transcription termination sites, alternative transcription start sites, multiple exon skipping, exon skipping, alternative 5′ and 3′ splice sites, and intron retention. **b,** Isoforms showing significant age-associated changes in usage. Each point represents an event detected for an isoform, colored by event class: alternative transcription start site (ATSS), alternative transcription termination site (ATTS), alternative 5′splice site (A5), exon skipping (ES), multiple exon skipping (MES), alternative 3′ splice site (A3), and intron retention (IR). Isoforms are labeled by gene symbol; individual isoforms may contain multiple event classes. **c,** Representative isoform switch in Macaca mulatta major histocompatibility complex class II, DM beta (MAMU-DMB). Transcript models show exon structure, annotated Pfam domains, predicted topology, subcellular localization, and coding status. Bar plots show gene-level expression, isoform-level expression, and isoform usage (isoform fraction, IF) by age group. **d,** AlphaFold-predicted structures of three MAMU-DMB proteoforms superimposed: ENSMMUT00000040899 (red), ENSMMUT00000040897 (blue), and ENSMMUT00000108347 (green). e, Representative isoform switch in bone gamma-carboxyglutamate protein (BGLAP), presented as in c. Six isoforms were detected, including five predicted coding isoforms and one predicted non-coding isoform. **f,** AlphaFold-predicted structures of two BGLAP proteoforms superimposed: ENSMMUT00000000123 (green) and ENSMMUT00000000124 (red). **g,** Representative isoform switch in LOC719702, presented as in c. **h,** AlphaFold-predicted structures of two LOC719702 proteoforms superimposed: ENSMMUT00000093320 (magenta; increased usage in aged animals) and ENSMMUT00000106706 (green). Isoform switches were identified using IsoformSwitchAnalyzeR (young, n = 6; aged, n = 5). In c, e, and g, bars show group means, and error bars indicate 95% confidence intervals. Statistical significance of isoform usage differences is indicated by *q < 0.05, ***q < 0.001, and ns, not significant.

AlphaFold models of ENSMMUT00000000123 (reference and ENSMMUT00000000124 are shown in Fig 9f. The reference model is a larger, predominantly helical protein, whereas ENSMMUT00000000124 is a much smaller protein composed of short helical segments and disordered regions. In the unannotated gene ENSMMUG00000050829 (LOC719702), total gene expression was unchanged with age, but usage of ENSMMUT00000106706 decreased from approximately 0.31 to 0.17 (q = 0.043) (Fig 9g). ENSMMUT00000093320 showed a non-significant increase, and ENSMMUT00000064737 was unchanged. All three isoforms are predicted to localize to the membrane and lysosome and contain a signal peptide, MHC_I, and C1-set domains, indicating an MHC class I-like protein. ENSMMUT00000064737 and ENSMMUT00000093320 share largely the same coding structure and differ mainly in their terminal exons. ENSMMUT00000106706 uses an alternative transcription start site and additionally contains a predicted transmembrane helix and an MHC_I_C cytoplasmic region. AlphaFold models of ENSMMUT00000093320 (reference) and ENSMMUT00000106706 are shown in Fig 9h. Both share an MHC-like fold of β-sheets and α-helices.

ENSMMUT00000106706 has an additional extended helical C-terminal segment, consistent with its transmembrane region. Age-related isoform remodelling in the hippocampus is therefore more extensive in females than in males, is driven predominantly by alternative transcription initiation and termination, and can either accompany a change in gene-level expression, as in BGLAP, or occur entirely without one, as in MAMU-DMB and LOC719702.

### Isoform co-expression modules capture age- and sex-specific transcriptional states and converge on Alzheimer’s disease risk isoforms

Having established that ageing remodels the hippocampal transcriptome at isoform resolution, we next asked whether these isoforms form coordinated co-expression networks with age in a sex-specific manner and whether such networks capture genetically defined AD risk. As above, we performed weighted gene co-expression network analysis (WGCNA) on variance-stabilized isoform expression. Hierarchical clustering of isoforms on topological overlap resolved discrete co-expression modules (Fig. 10a), whose composition is given in Supplementary Table 24.

**Fig 10.**
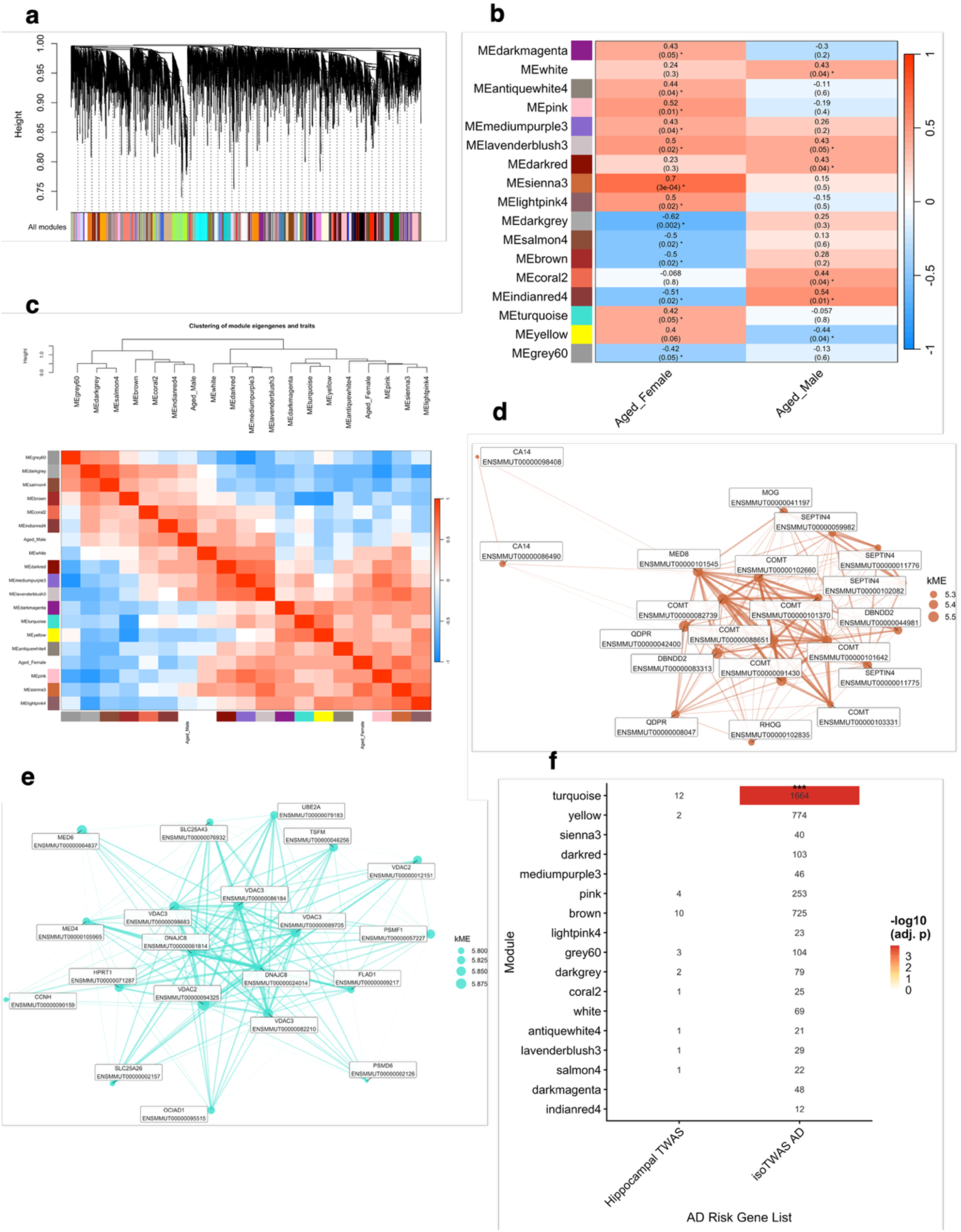
Isoform co-expression modules capture age- and sex-specific transcriptional states linked to AD. **a,**Hierarchical clustering of isoforms based on topological overlap identifies discrete co-expression modules, each labelled by a distinct color. Modules were defined using dynamic tree cutting, and only high-quality isoforms passing WGCNA quality control are shown. **b,** Heat map of Pearson correlations between module eigengenes and age–sex traits, highlighting modules significantly associated with aged females and aged males. The color scale denotes correlation strength and sign (red, positive; blue, negative), and each cell reports the correlation coefficient with the corresponding P value in parentheses; asterisks indicate significant associations (P < 0.05). **c,** Clustering of significant module eigengenes and traits with the corresponding eigengene adjacency heat map. The dendrogram summarizes eigengene similarity, and the adjacent heat map depicts pairwise eigengene correlations, where warm and cool colors indicate positively and negatively correlated eigengenes, respectively. **d,** Co-expression network for the sienna3 module. Nodes represent the top 20 isoforms ranked by module membership score (kME), scaled by kME value to reflect the strength of each isoform’s association with the module eigengene. Edges connect isoform pairs with strong topological overlap (TOM > 0.05), where edge width and transparency reflect the magnitude of co-expression. Node labels display the human ortholog gene name and corresponding rhesus macaque transcript ID. **e,** Co-expression network for the turquoise module outlined and interpreted as in **d**. **f,** Heatmap showing the overlap between significant rhesus macaque aging-associated co-expression modules (y-axis) and published Alzheimer’s disease risk gene lists (x-axis) derived from two transcriptome-wide association studies: 1. (Bhattacharya et al. 2023, Nature Genetics^54^) restricted to adult brain and Alzheimer’s disease associations, and 2. (Liu et al. 2021, PLOS Genetics^53^). Values within cells indicate the number of shared ortholog-mapped genes between each module and each AD risk gene list. Cell color represents the statistical significance of overlap expressed as −log10 Benjamini-Hochberg-adjusted P value. White cells indicate non-significant overlap; increasing color intensity indicates greater significance. Statistically significant overlaps are marked with asterisks (adjusted P < 0.05). Modules are ordered by significance and overlap across gene lists.

Module colors are assigned independently within each network; the modules described here are those of the isoform network and bear no relationship to the identically named modules of the gene-level network. Correlating module eigengenes with the age–sex traits revealed sex-specific structure (Fig. 10b). Ten modules were significantly associated with aging in females (P < 0.05). MEsienna3 showed by far the strongest positive association (r = 0.70, P = 3 × 10⁻⁴), followed by MEpink (r = 0.52), MElavenderblush3 and MElightpink4 (r = 0.50 each), MEantiquewhite4 (r = 0.44), MEdarkmagenta and MEmediumpurple3 (r = 0.43 each). MEturquoise (r = 0.42); MEdarkgrey (r = −0.62), MEindianred4 (r = −0.51), MEsalmon4 and MEbrown (r = −0.50 each) and MEgrey60 (r = −0.42) were negatively correlated with age in females. Fewer modules reached significance in males, and they were largely distinct: MEindianred4 (r = 0.54), MEcoral2 (r = 0.44), MElavenderblush3 and MEdarkred (r = 0.43 each) and MEwhite (r = 0.41) were positively correlated with aging, and MEyellow (r = −0.44) negatively. Only MElavenderblush3 was positively associated in both sexes, whereas MEindianred4 was negatively associated in females and positively in males, and several strongly female-associated modules, including MEsienna3, showed no male association at all (Fig 10b).

Hierarchical clustering of the significant eigengenes together with the traits confirmed that this divergence is a property of the network as a whole rather than of individual modules (Fig. 10c). The aged-male trait clustered with MEgrey60, MEdarkgrey, MEsalmon4, MEbrown, MEcoral2 and MEindianred4, while the aged-female trait clustered with a separate branch containing MEantiquewhite4, MEpink, MEsienna3 and MElightpink4, with the eigengene adjacency heat map showing these two groups to be mutually anticorrelated (Fig 10c). Age-related isoform co-expression is therefore organized into several co-regulated groups that separate by sex rather than into a single unified aging program.

To characterize the two most informative female-associated modules, we visualized their top 20 isoforms ranked by module membership (kME). The sienna3 network, the most strongly female-associated module, was densely interconnected and built predominantly from oligodendrocyte and myelin-associated transcripts, with multiple independent isoforms of *COMT* and *SEPTIN4* occupying the most central positions alongside *MOG, QDPR, DBNDD2, CA14, MED8* and *RHOG* (Fig. 10d). The turquoise network was similarly dense but functionally distinct, comprising mitochondrial and proteostatic transcripts: multiple isoforms of *VDAC2* and *VDAC3* and of *DNAJC8* formed the core, together with *MED4, MED6, SLC25A26, SLC25A43, TSFM, UBE2A, PSMD6, PSMF1, FLAD1, HPRT1, CCNH* and *OCIAD1* (Fig. 10e).

Finally, we asked whether any module is enriched for genetically implicated AD risk genes. To assess this, we tested each module against two publicly available lists by hypergeometric test with Benjamini–Hochberg correction across all module-by-list comparisons (Fig. 10f). Analysis gainst the hippocampal TWAS list of 43 genes described by Liu et.al^53^, revealed no module significant overlap after correction; the largest raw overlaps were with MEblue (13 genes, including APOE, BIN1, and CD55; P = 0.013) and MEred (7 genes, P = 0.015), neither surviving adjustment (supplementary table 25). Against the isoTWAS AD list described by Bhattacharya et al^54^, one module was significantly enriched: MEturquoise shared 1,664 of its 2,401 ortholog-mapped genes with the risk list (P = 1.4 × 10⁻⁵, adjusted P = 1.2 × 10⁻⁴, Jaccard index 0.27), the only significant overlap in either comparison. MEyellow (774 shared genes, P = 0.0024) and MEgreenyellow (180 genes, P = 0.012) were nominally enriched but did not survive correction (Fig 10f; Supplementary Table 25). MEturquoise is a female-associated module (r = 0.42, P = 0.05) whose hub isoforms are mitochondrial carriers, chaperones and proteasome subunits (Fig. 10e), and the shared gene set includes the proteasome complement (*PSMA2–PSMA4, PSMB3, PSMC5, PSMC6, PSMD6, PSMD13, PSMD14, PSME1, PSMF1*), mitochondrial respiratory and import machinery (*NDUFS1, NDUFS2, NDUFA5, NDUFA10, SDHB, TIMM22, SAMM50, VDAC2*), and established AD-associated genes including *TREM2, INPP5D, APH1A, SNCA, UCHL1, MAPT*, *LRRC37A/B, PTK2B*, *CD55* and *APOC2* (Supplementary table 26). This is consistent with the original isoTWAS analysis^54^, in which AD-prioritized genes were themselves enriched for proteasome regulation, and it links the isoform network that changes with age in female macaques to the isoform-level genetic architecture of human AD. Together, these findings highlight the value of isoform-resolved analysis for capturing the transcriptional complexity of brain ageing and revealing molecular changes that may be obscured by gene-level approaches.

## Discussion

In this study, we leveraged long-read RNA sequencing to characterize age-associated transcriptomic remodeling in the rhesus macaque hippocampus at both gene and isoform resolution. We show that ageing is associated with transcriptional changes that occur at the gene level and at the level of alternative splicing among genes relevant to neurodegenerative susceptibility, neuroinflammation, and oxidative phosphorylation. Although both sexes exhibited enrichment of core neuroinflammatory pathways, including interferon, TNFα/NF-κB and IL6/JAK/STAT3 signalling, the genes driving these responses differed substantially between males and females. Co-expression analysis similarly revealed distinct ageing-associated network states, with female ageing characterized by mitochondrial, synaptic, senescence and proteostasis-related changes, whereas male ageing showed stronger oligodendrocyte and myelin-associated remodeling. Thus, neuroinflammation emerges as a shared feature of hippocampal ageing, but its transcriptional context is strongly sex dependent.

This distinction may be relevant to the differential susceptibility to neurodegenerative disease observed across the lifespan. Estrogen signalling is particularly relevant to female brain ageing because estrogen receptors are widely expressed in the hippocampus, where estrogen regulates synaptic plasticity^55^, mitochondrial function^46^, oxidative stress^56^, and glial signalling^57^.

Experimental and translational studies further implicate estrogen in the regulation of microglial and astrocytic inflammatory responses and in processes relevant to AD pathology. Loss of ovarian estrogen during menopause may therefore alter the molecular environment in which age-related inflammation develops, although the contribution of estrogen to AD susceptibility is complex and clinical evidence for hormone therapy and dementia prevention remains mixed. Our data do not establish hormone loss as the basis of the female-specific ageing trajectory, but the convergence of hormone-responsive, mitochondrial, synaptic, and inflammatory pathways provides a molecular context in which endocrine changes could influence hippocampal resilience during ageing. Cross-species network analysis identified two conserved programs associated with both normative macaque ageing and the human AD hippocampus. A 553-gene oxidative-phosphorylation/proteasome network decreased with age in females and was also reduced in human AD, whereas an oligodendrocyte/myelin program increased with age in males and was represented within an AD-associated human module. These findings do not indicate that normative macaque ageing recapitulates AD, but rather that distinct molecular trajectories engaged during ageing converge on transcriptional programs altered in human disease. This provides a framework for using the macaque model to investigate how age-associated changes in mitochondrial, proteostatic, and glial biology may create molecular states relevant to neurodegenerative vulnerability. A major implication of this study is that many age-associated changes are incompletely captured at the gene level. Many differentially expressed isoforms clearly separated young and aged animals despite little or no change in total expression of their parent genes, while differential transcript usage identified switching events driven predominantly by alternative transcript start and end sites. These findings demonstrate that ageing can remodel transcript composition without substantially altering overall gene abundance. Long-read sequencing therefore, reveals a regulatory layer that is obscured when transcripts are aggregated to genes, providing a more complete view of the molecular complexity of brain ageing.

The translational relevance of this isoform-level remodeling was supported by its relationship to human AD. A female-associated isoform module was enriched for genes prioritized by an AD isoTWAS, including mitochondrial carriers, chaperones, and proteasome components. This overlap links an age-associated female hippocampal network with genes implicated in AD susceptibility and suggests that isoform-level regulation may provide an important molecular interface between brain ageing and genetic risk for neurodegeneration. Conversely, the smaller hippocampal TWAS set of 43 genes showed no significant module enrichment, which likely reflects limited statistical power. These findings highlight the value of resolving transcript-level regulation when investigating how genetic risk is expressed within the ageing brain. Together, these results suggest that aging engages divergent immune-regulatory trajectories in males and females, potentially contributing to sex differences in neurodegenerative disease susceptibility and progression. Crucially, our use of long-read sequencing enabled transcript-level analyses that extend beyond gene abundance and capture changes in isoform composition that would otherwise remain undetected. While short-read RNA sequencing fragments transcripts and can obscure full-length isoform structure, long-read sequencing provides direct resolution of alternatively spliced transcripts, allowing for better quantification of isoform usage. This is particularly important in the aging brain, where subtle shifts in splicing patterns may have outsized functional consequences. Importantly, our results reveal that age-associated changes occur independently of differential gene expression. This suggests that aging not only alters which genes are expressed, but also reshapes the relative usage of distinct transcripts encoded by those genes. Such isoform remodeling has the potential to influence protein structure, localization, interaction partners, and aggregation propensity, thereby modulating cellular function without changing total transcript abundance. In the context of neurodegeneration, this distinction is critical, as small changes in isoform balance may influence proteoform ratios and can influence function depending on the protein’s structure. Taken together, our findings support a model in which aging establishes a sex-specific transcriptomic landscape characterized by both shared inflammatory signatures and divergent regulatory programs. This sex bias is consistent with epidemiological data showing differential risk, prevalence, and progression of neurodegenerative diseases between men and women^58,59^, and suggests that transcriptomic regulation, including isoform usage, may contribute to these differences at a molecular level. Several limitations should be acknowledged. While transcriptomic changes provide insight into regulatory processes, future studies integrating proteomic analyses and functional assays will be necessary to establish causal links between isoform remodeling and cellular phenotypes. In addition, the hippocampus comprises multiple neuronal and glial cell types, and bulk sequencing cannot fully resolve cell-type-specific contributions. Single-cell or spatial long-read transcriptomic approaches will be essential to define the cellular origins of these aging-associated isoform programs. Despite these limitations, this study demonstrates the power of long-read transcriptomics to uncover regulatory features of brain aging that are invisible to gene-centric analyses alone. By revealing sex-biased gene- and isoform-level remodeling in pathways central to inflammation and neurodegeneration, our work highlights alternative splicing as a component of brain inflammaging and a potential contributor to sex-specific vulnerability to age-associated neurodegenerative disease. These findings underscore the importance of incorporating isoform-aware strategies into studies of aging and suggest that targeting splicing regulation may offer new opportunities for sex-informed therapeutic intervention.

## Methods

Sample Collection Hippocampal samples were selected from the tissue archive of the Aging Primate Resource at the Oregon National Primate Research Center. These samples originated from the tissue distribution program and the process followed the guidelines from the 2013 American Veterinary Medical Associationfor the Euthanasia of Animals. Each brain was flushed with saline and the right hippocampus was removed, dissected into thirds in the coronal plane, frozen in liquid nitrogen and archived at −80 C. The middle third of each sample was used and contained the entirety of the hippocampus. Samples were all from animals that had no evidence of abnormal behavior or gross lesions in the brain.

### RNA Extraction

RNA was extracted from frozen hippocampal tissues as previously described (PMID: 40824352). Briefly, automated nucleic acid extractions were performed on pulverized and homogenized samples using the Qiagen AllPrep DNA/RNA/miRNA Universal Kit on a QIAcube by the ONPRC Primate Genetics Core. RNA integrity and concentration were assessed using an Agilent Bioanalyzer Qubit fluorometry. Aliquots were frozen at −80C prior to cDNA synthesis and library preparation.

### Oxford Nanopore cDNA Barcoding Library Preparation and Sequencing

Hippocampal RNA concentration was quantified using the Qubit RNA High Sensitivity (HS) Assay Kit (Thermo Fisher Scientific, Q32852). Libraries were prepared following the manufacturer’s protocol for the Oxford Nanopore cDNA Barcoding Kit (SQK-PCB114.24). Briefly, total hippocampal RNA was normalized across samples and reverse transcribed into cDNA using Maxima H Minus Reverse Transcriptase (200 U/μl) (ThermoFisher, EP0752), enabling strand switching and incorporation of unique molecular identifiers (UMIs). The resulting cDNA was amplified by PCR using LongAmp Hot Start Taq 2X Master Mix (NEB, M0533), which facilitated the attachment of sample-specific barcode primers. PCR product concentration was determined using the Qubit dsDNA HS Assay Kit (Thermo Fisher Scientific, Q32851), and fragment size distribution was evaluated using the Agilent D5000 ScreenTape Assay (5067-5588, 5067-5589, 5067-5590). Equimolar quantities of barcoded PCR products were pooled, and sequencing adapters were ligated to generate the final library. The library was sequenced on four R10.4.1 PromethION flow cells (Oxford Nanopore, FLO-PRO114M) using the PromethION 2 Solo (P2 Solo) (Oxford Nanopore, PRO-SEQ002).

### Data Preprocessing and Analysis Long-Read RNA Sequencing

Raw POD5 files were basecalled using Dorado (https://github.com/nanoporetech/dorado) to generate FASTQ files. Low-quality reads were filtered using Chopper (https://github.com/wdecoster/chopper). Processed FASTQ files were then analyzed using the Oxford Nanopore wf-transcriptomes pipeline (https://github.com/epi2me-labs/wf-transcriptomes). Briefly, reads were aligned to the reference transcriptome and genome using minimap2 (https://github.com/lh3/minimap2). Raw long-read transcript counts were obtained after running the Oxford Nanopore Technologies (ONT) WF-Transcriptomes pipeline using the Macaca_mulatta.Mmul_10.dna.toplevel.fa reference genome, the Macaca_mulatta.Mmul_10.cdna.all.fa reference transcriptome, and the Macaca_mulatta.Mmul_10.115.gtf reference annotation. Gene-level and transcript-level counts were generated using the ONT wf-transcriptomes pipeline. The resulting count matrices were imported into R and filtered to remove non-informative columns, with gene and transcript Ensembl identifiers assigned as row names. Sample metadata included variables for sex, age, and batch, which were encoded as categorical factors. Metadata and sample ids used in figures can be found in Supplementary Tables 13 and 14.

### Differential Gene and Isoform Expression

Gene and isoform level differential expression analyses were performed using the DESeq2 package (v1.40.2)^60^ in R (v4.3.1). DESeq2 normalization and model fitting were performed using the default negative binomial framework implemented in the DESeq() function as previously described^60^. Normalized counts generated by DESeq2 were extracted and saved for downstream analyses and visualization. For sex-stratified analyses, samples were analyzed separately by sex using the design formula ∼ batch + age, with batch included to account for technical variation and age coded as young versus aged. Female and male age-associated expression changes were therefore estimated independently within each sex. Statistical significance was defined as a Benjamini–Hochberg adjusted P value ≤ 0.05. DESeq2-normalized counts were used for downstream analyses. For visualization, normalized counts and variance-stabilized expression values were batch-corrected using limma::removeBatchEffect, with age retained in the design matrix. Significant genes were annotated by mapping Ensembl gene identifiers to gene symbols using a species-specific annotation table.

### Weighted Gene Co-expression Network Analysis (WGCNA)

Weighted gene co-expression network analysis (WGCNA) was performed on variance-stabilized isoform expression to identify co-expression modules associated with sex-specific ageing.

Isoforms were first filtered by retaining those with ≥10 counts in at least six samples, corresponding to the smallest experimental group. Variance-stabilizing transformation was then performed using DESeq2 (blind = FALSE), and the upper 75% most variable isoforms were retained based on expression variance.

To reduce technical variation associated with differences in transcript detection, the detection rate was assessed across samples and confirmed not to be significantly associated with age, sex, or their interaction. Detection rate was subsequently included as a covariate during adjustment of the VST expression matrix using limma::removeBatchEffect as previously described^61^, while the age-by-sex design was retained to preserve biological variation of interest. Surrogate variable analysis (SVA) was then performed on the adjusted matrix using SVA version 3.60.0^62^, and four surrogate variables were removed while preserving the age-by-sex design. The resulting expression matrix was transposed to generate a samples-by-isoforms matrix for WGCNA.

Signed co-expression networks were constructed using blockwiseModules, with a minimum module size of 30 isoforms and a maximum block size of 25,000 isoforms. The soft-thresholding power was selected using pickSoftThreshold based on scale-free topology and network connectivity. Modules were identified by topological overlap-based hierarchical clustering and dynamic tree cutting, and modules with highly correlated eigengenes were merged using a cut height of 0.10. Module eigengenes were calculated as the first principal component of each module. Module–trait relationships were assessed by correlating module eigengenes with aged-female and aged-male traits, with statistical significance defined as P < 0.05. Signed module membership (kME) was calculated for each isoform to characterize connectivity within its assigned module. Module gene membership can be found in Supplementary Table 15.

### Module–Trait Relationships

Module eigengenes were correlated with age- and sex-related traits using Pearson correlation. Age and sex were encoded as binary variables, and additional interaction traits were generated to capture age effects within each sex (Age_Female and Age_Male). Statistical significance of module–trait correlations was assessed using Student asymptotic p-values. Modules showing significant associations (p < 0.05) with at least one trait were retained for downstream analyses. Module–trait relationships were visualized using labeled heatmaps, with correlation coefficients and corresponding p-values displayed. Significant associations were denoted with an asterisk.

### Eigengene Network and Module Visualization

To examine relationships among significant modules and traits, eigengene adjacency networks were constructed by correlating significant module eigengenes with trait variables, followed by hierarchical clustering based on eigengene dissimilarity. These relationships were visualized using combined dendrogram and heatmap representations. Gene-level dendrograms were generated to visualize the hierarchical clustering of isoforms within significant modules. Dendrograms highlighting only genes belonging to significant modules were produced to emphasize age- and sex-associated co-expression structure.

### Differential Transcript Usage (DTU) Analysis

Differential transcript usage was assessed using the IsoformSwitchAnalyzeR R package version^62^, with statistical testing performed using its DEXSeq-based isoform switch analysis. Transcript-level count matrices, sample metadata, the Macaca mulatta GTF annotation, and cDNA FASTA sequences were imported using the importRdata function. Analyses were performed separately for females and males, with age as the primary condition and batch included as a covariate where applicable. Transcripts and genes with low expression were removed using the prefilter function, requiring a minimum isoform expression of 3 and gene expression of 5, and genes containing only a single isoform were excluded. Differential isoform usage between young and aged animals was then evaluated using isoformSwitchTestDEXSeq(), which applies a DEXSeq-based model to identify changes in isoform usage between conditions. Statistical significance was defined as a false discovery rate (FDR) < 0.05 together with an absolute change in isoform fraction (|ΔIF|) ≥ 0.05. Analyses were restricted to genes with significant isoform switching using reduceToSwitchingGenes = TRUE, while retaining all isoforms for subsequent characterization. Significant switching genes were extracted using extractTopSwitches. Alternative splicing events among switching isoforms were subsequently characterized using analyzeAlternativeSplicing, including exon skipping, intron retention, alternative 5′ and 3′ splice sites, and alternative transcription start and termination sites.

### Pathway Analysis of WGCNA Modules

Genes assigned to WGCNA modules were assessed for functional enrichment using the clusterProfiler^63^ and msigdbr R packages. Rhesus macaque genes were mapped to human orthologs using biomaRt, and the resulting human gene symbols were used for downstream enrichment analysis. Gene set enrichment was performed using the clusterProfiler::enricher function against MSigDB gene sets, including Hallmark (H), C2, and Gene Ontology (C5) collections. All ortholog-mapped genes represented in the WGCNA dataset were used as the background gene universe to account for the genes available for network analysis. Enrichment was considered significant at an adjusted P value < 0.05, with gene sets required to contain at least three genes and no more than 500 genes. Significant pathways were ranked by adjusted P value, and the top enriched pathways were visualized using bar or dot plots, with bar length representing the number of module genes contributing to each pathway and color indicating statistical significance.

### Gene Set Enrichment Analysis

Gene set enrichment analysis (GSEA) was performed using the Gene Set Enrichment Analysis (GSEA v4.3.3) software (Broad Institute, Cambridge, MA) to identify pathways differentially enriched across our experimental groups. The gene-level normalized count matrix served as input, along with a metadata file classifying each sample into one of four experimental groups: young female, young male, aged female, and aged male. The Hallmark gene set collection (h.all.v2025.1.Hs.symbols.gmt) from the Molecular Signatures Database (MSigDB v2025.1) was used as the reference database. The analysis was conducted using 1,000 phenotype permutations, a weighted enrichment statistic, and signal-to-noise ratio as the gene-ranking metric. Gene sets containing fewer than 15 or more than 1000 genes were excluded to reduce bias from small or overly broad categories. Normalization of enrichment scores was performed using the meandiv method. The top 20 significantly enriched gene sets were selected for visualization based on normalized enrichment score (NES) and false discovery rate (FDR) q-value < 0.05.

### Cross Species Shared Gene Analysis

Cross-species network analysis was performed using weighted gene co-expression network analysis (WGCNA) module assignments generated independently for rhesus macaque and human hippocampus. Rhesus modules with an age-associated module–trait P value < 0.05 in either sex were retained, and human modules with a minimum module–trait P value < 0.05 across the available clinical groups were similarly selected. Rhesus macaque genes were mapped to human orthologs using Ensembl BioMart, and genes without a corresponding human ortholog represented in the human dataset were excluded. For analyses requiring a common gene namespace, orthologous genes were represented using human gene symbols. Cross-species module overlap was assessed using shared orthologous genes between significant rhesus and human modules. Pairwise module overlap was quantified using the Jaccard index and tested using one-sided hypergeometric and Fisher’s exact tests, with the union of genes represented across the tested modules defining the background universe. P values were adjusted for multiple comparisons using the Benjamini–Hochberg method, with an adjusted P value < 0.05 considered significant.

### Alzheimer’s disease risk gene overlap analysis

AD-associated gene lists were compiled from an isoform-level transcriptome-wide association study (isoTWAS), and a hippocampal TWAS. For isoTWAS, results were restricted to Alzheimer’s disease (ALZ), the Adult dataset and the isoTWAS method, followed by filtering for permutation P < 0.05. Hippocampal TWAS genes were filtered at P < 0.05. For each module and AD-associated gene list, the number of overlapping genes was determined and statistical significance was assessed using a one-sided hypergeometric test. The background universe comprised all unique genes represented in the annotated WGCNA dataset. P values were adjusted independently within each AD-associated gene list using the Benjamini–Hochberg procedure, with an adjusted P value < 0.05 considered significant. Gene-set similarity was additionally quantified using the Jaccard index, calculated as the number of shared genes divided by the union of module and AD-associated genes. Significant overlaps were visualized using ggplot2.

### Protein Modeling

Protein structures were predicted by AlphaFold v6. Structures were superimposed in using the Matchmaker tool in UCSF Chimerax v1.11, with Needleman-Wunsch algorithm and BLOSUM-62 matrix for sequence alignment.

### Data Output and Reproducibility

Raw and processed data will be available on GEO and the code will be published in a github repository.

### Statistical information

Module–trait associations were assessed using Pearson correlation between module eigengenes and binary age-by-sex traits. Correlation P values were calculated using the Student’s t-distribution, with P < 0.05 considered significant. Modules with a significant association with either aged-female or aged-male traits were retained for downstream analysis. Pathway enrichment was assessed using one-sided hypergeometric over-representation tests, with P values adjusted for multiple comparisons using the Benjamini–Hochberg (BH) method. Pathways with adjusted P < 0.20 were retained. Cross-species module overlap was evaluated using one-sided hypergeometric and Fisher’s exact tests, with BH-adjusted P < 0.05 considered significant. The Jaccard index was used to quantify gene-set similarity. AD-associated gene overlap was similarly assessed using one-sided hypergeometric tests with BH-adjusted P < 0.05 considered significant. Each animal was treated as one independent biological replicate. The gene-level dataset (illumina short-read sequencing) included n = 6 per age-by-sex group, while the long-read dataset included n = 6 young females, 5 aged females, 6 young males, and 5 aged males, after excluding two animals that were not well-resolved during de-multiplexing.

## Data Availability

All sequencing data will be made available upon publication via GEO.

## Code Availability

We did not develop new code for this study. Software tools and code used in this study are publicly available and described in the Methods section.

## Acknowledgements

The authors would like to thank Drs. R. Keith Reeves and C. Sabrina Tan for helpful discussions. This research was funded in part by the National Institutes of Health (United States) 7K01OD031899-03 and 3K01OD031899-03S1 (to D.R.R). Services reported in this publication provided by the NHP Aging Resource were supported by the National Institute on Aging and the Office of the Director, National Institutes of Health under Award Number P51OD011092 and used the services of the ONPRC Primate Genetics Core (RRID:SCR_027583). The content is solely the responsibility of the authors and does not necessarily represent the official views of the National Institutes of Health.

## Ethics declarations

The authors declare no competing interests.

## Supplementary Information

- Supplementary Figures 1-12
- Supplementary Tables 1-14
- Supplementary Tables 15-26

## Contributions

S.K. and D.R.R. conceived the work. V.H.A., S.P. and S.S. carried out experiments. V.H.A, S.P. S.K. and D.R.R. analyzed the data. V.H.A wrote the initial draft of the manuscript. V.H.A, S.P., S.S., S.K. and D.R.R. reviewed and approved the final text.

## References

1. Troen, B. R. The biology of aging. Mt Sinai J Med 70, 3–22 (2003).

2. Xi, J.-Y. et al. Effects of population aging on quality of life and disease burden: a population-based study. glob health res policy 10, 2 (2025).

3. Shreya, D., Fish, P. N. & Du, D. Navigating the Future of Elderly Healthcare: A Comprehensive Analysis of Aging Populations and Mortality Trends Using National Inpatient Sample (NIS) Data (2010-2024). Cureus 17, e80442.

4. Lampropoulos, I. C., Malli, F., Sinani, O., Gourgoulianis, K. I. & Xiromerisiou, G. Worldwide trends in mortality related to Parkinson’s disease in the period of 1994–2019: Analysis of vital registration data from the WHO Mortality Database. Front. Neurol. 13, (2022).

5. Bellenguez, C. et al. New insights into the genetic etiology of Alzheimer’s disease and related dementias. Nat Genet 54, 412–436 (2022).

6. Polymeropoulos, M. H. et al. Mutation in the α-Synuclein Gene Identified in Families with Parkinson’s Disease. Science 276, 2045–2047 (1997).

7. Soraci, L. et al. Neuroinflammaging: A Tight Line Between Normal Aging and Age-Related Neurodegenerative Disorders. Aging Dis 15, 1726–1747 (2024).

8. Deng, X.-H., Bertini, G., Xu, Y.-Z., Yan, Z. & Bentivoglio, M. Cytokine-induced activation of glial cells in the mouse brain is enhanced at an advanced age. Neuroscience 141, 645–661 (2006).

9. Elobeid, A., Libard, S., Leino, M., Popova, S. N. & Alafuzoff, I. Altered Proteins in the Aging Brain. J Neuropathol Exp Neurol 75, 316–325 (2016).

10. Cumplido-Mayoral, I. et al. Neuroimaging-derived biological brain age and its associations with glial reactivity and synaptic dysfunction cerebrospinal fluid biomarkers. Mol Psychiatry 30, 3718–3728 (2025).

11. Sartori, A. C., Vance, D. E., Slater, L. Z. & Crowe, M. The Impact of Inflammation on Cognitive Function in Older Adults: Implications for Health Care Practice and Research. J Neurosci Nurs 44, 206–217 (2012).

12. Arosio, B. et al. Inflammaging and the sex-frailty paradox. Aging Clin Exp Res 37, 282 (2025).

13. Jin, K. et al. Brain-wide cell-type-specific transcriptomic signatures of healthy ageing in mice. Nature 638, 182–196 (2025).

14. Merkin, J., Russell, C., Chen, P. & Burge, C. B. Evolutionary Dynamics of Gene and Isoform Regulation in Mammalian Tissues. Science 338, 1593–1599 (2012).

15. Lerch, J. K., Bixby, J. L. & Lemmon, V. P. Isoform diversity and its importance for axon regeneration. Neuropathology 32, 420–431 (2012).

16. Patowary, A. et al. Developmental isoform diversity in the human neocortex informs neuropsychiatric risk mechanisms. Science 384, eadh7688 (2024).

17. Beyer, K. α-Synuclein structure, posttranslational modification and alternative splicing as aggregation enhancers. Acta Neuropathol 112, 237–251 (2006).

18. Hoyer, W., Cherny, D., Subramaniam, V. & Jovin, T. M. Impact of the Acidic C-Terminal Region Comprising Amino Acids 109−140 on α-Synuclein Aggregation in Vitro. Biochemistry 43, 16233–16242 (2004).

19. Bhadra, M., Howell, P., Dutta, S., Heintz, C. & Mair, W. B. Alternative splicing in aging and longevity. Hum Genet 139, 357–369 (2020).

20. Wang, K. et al. Comprehensive map of age-associated splicing changes across human tissues and their contributions to age-associated diseases. Sci Rep 8, 10929 (2018).

21. Mazin, P. et al. Widespread splicing changes in human brain development and aging. Mol Syst Biol 9, 633 (2013).

22. Floor, S. N. & Doudna, J. A. Tunable protein synthesis by transcript isoforms in human cells. eLife 5, e10921.

23. Ueda, M. T. et al. Functional and dynamic profiling of transcript isoforms reveals essential roles of alternative splicing in interferon response. Cell Genomics 4, (2024).

24. Lee, F. F. Y. & Alper, S. Alternative pre-mRNA splicing as a mechanism for terminating Toll-like Receptor signaling. Front. Immunol. 13, (2022).

25. Robinson, E. K. et al. Inflammation drives alternative first exon usage to regulate immune genes including a novel iron-regulated isoform of Aim2. eLife 10, e69431 (2021).

26. Bettio, L. E. B., Rajendran, L. & Gil-Mohapel, J. The effects of aging in the hippocampus and cognitive decline. Neuroscience & Biobehavioral Reviews 79, 66–86 (2017).

27. Buss, E. W. et al. Cognitive aging is associated with redistribution of synaptic weights in the hippocampus. Proceedings of the National Academy of Sciences 118, e1921481118 (2021).

28. Wearn, A. et al. Quantitative MRI of the hippocampus reveals microstructural trajectories of aging and Alzheimer’s disease pathology. Proceedings of the National Academy of Sciences 122, e2502674122 (2025).

29. Wu, Y. et al. Multimodal transcriptomics reveal neurogenic aging trajectories and age-related regional inflammation in the dentate gyrus. Nat Neurosci 28, 415–430 (2025).

30. Femminella, G. D. et al. Does Microglial Activation Influence Hippocampal Volume and Neuronal Function in Alzheimer’s Disease and Parkinson’s Disease Dementia? Journal of Alzheimer’s Disease 51, 1275–1289 (2016).

31. Spangenberg, E. E. et al. Eliminating microglia in Alzheimer’s mice prevents neuronal loss without modulating amyloid-β pathology. Brain 139, 1265–1281 (2016).

32. Gillies, G. E., Pienaar, I. S., Vohra, S. & Qamhawi, Z. Sex differences in Parkinson’s disease. Front Neuroendocrinol 35, 370–384 (2014).

33. Kulminski, A. M. et al. Sex-specific genetic susceptibility to Alzheimer’s disease and diabetes. Alzheimers Dement 21, e103096 (2025).

34. Olivieri, F. et al. Sex/gender-related differences in inflammaging. Mechanisms of Ageing and Development 211, 111792 (2023).

35. Lu, Y. et al. Macaque Brainnetome Atlas: A multifaceted brain map with parcellation, connection, and histology. Science Bulletin 69, 2241–2259 (2024).

36. Downs, J. L. & Urbanski, H. F. Neuroendocrine Changes in the Aging Reproductive Axis of Female Rhesus Macaques (Macaca mulatta)1. Biol Reprod 75, 539–546 (2006).

37. Messaoudi, I., Estep, R., Robinson, B. & Wong, S. W. Nonhuman Primate Models of Human Immunology. Antioxid Redox Signal 14, 261–273 (2011).

38. Gibbs, R. A. et al. Evolutionary and Biomedical Insights from the Rhesus Macaque Genome. Science 316, 222–234 (2007).

39. Uno, H. The incidence of senile plaques and multiple infarction in aged Macaque brain. Neurobiology of Aging 14, 673–674 (1993).

40. Heilbroner, P. L. & Kemper, T. L. The cytoarchitectonic distribution of senile plaques in three aged monkeys. Acta Neuropathol 81, 60–65 (1990).

41. Anderson, T. J. et al. Patterns of transcriptomic aging in the hippocampus of rhesus macaques highlight midlife transitions. GeroScience 48, 2913–2929 (2026).

42. Dias, K., Lemoine, S., Thomas-Chollier, M., Le Crom, S. & Jourdren, L. ToulligQC 2: fast and comprehensive quality control for Oxford Nanoporesequencing data. (2022).

43. Yuan, S. et al. CHIT1 regulates the neuroinflammation and phagocytosis of microglia and suppresses Aβ plaque deposition in Alzheimer’s disease. The Journal of Pathology 265, 330–341 (2025).

44. Subramanian, A. et al. Gene set enrichment analysis: A knowledge-based approach for interpreting genome-wide expression profiles. Proceedings of the National Academy of Sciences 102, 15545–15550 (2005).

45. Spence, R. D. et al. Estrogen Mediates Neuroprotection and Anti-Inflammatory Effects during EAE through ERα Signaling on Astrocytes But Not through ERβ Signaling on Astrocytes or Neurons. J. Neurosci. 33, 10924–10933 (2013).

46. Yue, X. et al. Brain estrogen deficiency accelerates Aβ plaque formation in an Alzheimer’s disease animal model. Proc Natl Acad Sci U S A 102, 19198–19203 (2005).

47. Appleman, M.-L. et al. Effect of hormone replacement therapy on amyloid beta (Aβ) plaque density in the rhesus macaque amygdala. Front. Aging Neurosci. 15, (2024).

48. Zuhlsdorff, K. et al. Emotional and cognitive effects of menopause and hormone replacement therapy. Psychological Medicine 56, e24 (2026).

49. Li, R., Cui, J. & Shen, Y. Brain Sex Matters: estrogen in cognition and Alzheimer’s disease. Mol Cell Endocrinol 389, 13–21 (2014).

50. Briceno Silva, G., et al. Influence of the Onset of Menopause on the Risk of Developing Alzheimer’s Disease. Cureus 16, e69124.

51. Disouky, A. et al. Human hippocampal neurogenesis in adulthood, ageing and Alzheimer’s disease. Nature 652, 1264–1273 (2026).

52. Bethea, C. L., Kohama, S. G., Reddy, A. P. & Urbanski, H. F. Ovarian steroids regulate gene expression in the dorsal raphe of old female macaques. Neurobiology of Aging 37, 179–191 (2016).

53. Liu, N. et al. Hippocampal transcriptome-wide association study and neurobiological pathway analysis for Alzheimer’s disease. PLOS Genetics 17, e1009363 (2021).

54. Bhattacharya, A. et al. Isoform-level transcriptome-wide association uncovers genetic risk mechanisms for neuropsychiatric disorders in the human brain. Nat Genet 55, 2117–2128 (2023).

55. Liu, F. et al. Activation of estrogen receptor-β regulates hippocampal synaptic plasticity and improves memory. Nat Neurosci 11, 334–343 (2008).

56. Zhang, Q.-G. et al. Estrogen Attenuates Ischemic Oxidative Damage via an Estrogen Receptor α-Mediated Inhibition of NADPH Oxidase Activation. J Neurosci 29, 13823–13836 (2009).

57. Goenaga, J. et al. Estradiol Mediates Astrocyte-Neuron Communication in the Hippocampus. Mol Neurobiol 62, 10468–10483 (2025).

58. Stoccoro, A. Epigenetic Mechanisms Underlying Sex Differences in Neurodegenerative Diseases. Biology 14, 98 (2025).

59. Bianco, A., Antonacci, Y. & Liguori, M. Sex and Gender Differences in Neurodegenerative Diseases: Challenges for Therapeutic Opportunities. International Journal of Molecular Sciences 24, 6354 (2023).

60. Love, M. I., Huber, W. & Anders, S. Moderated estimation of fold change and dispersion for RNA-seq data with DESeq2. Genome Biol 15, 550 (2014).

61. Ritchie, M. E. et al. limma powers differential expression analyses for RNA-sequencing and microarray studies. Nucleic Acids Res 43, e47 (2015).

62. Vitting-Seerup, K. & Sandelin, A. IsoformSwitchAnalyzeR: analysis of changes in genome-wide patterns of alternative splicing and its functional consequences. Bioinformatics 35, 4469–4471 (2019).

63. Yu, G., Wang, L.-G., Han, Y. & He, Q.-Y. clusterProfiler: an R Package for Comparing Biological Themes Among Gene Clusters. OMICS 16, 284–287 (2012).

64. Leek, J. T., et al. sva: Surrogate Variable Analysis. R package version 3.56.0. Bioconductor 10.18129/B9.bioc.sva (2025)

