## Supplementary Figures 1-12 for "Sex-specific gene and isoform expression characterizes hippocampal aging in rhesus macaques"

### Author Information


#### Affiliations

**Burnett School of Biomedical Sciences, University of Central Florida, Orlando, Florida, United States of America**

Victoria Harper-Alexander-1, Saida Sadikova-3 & Daniel Ramos Ram-5

**Oregon National Primate Research Center, Beaverton, Oregon, United States of America**

Sam Peterson-2 & Steven Kohama-4

#### Corresponding author

Correspondence to: Author-5

Corresponding author ORCID ID: 0000-0002-9841-9525

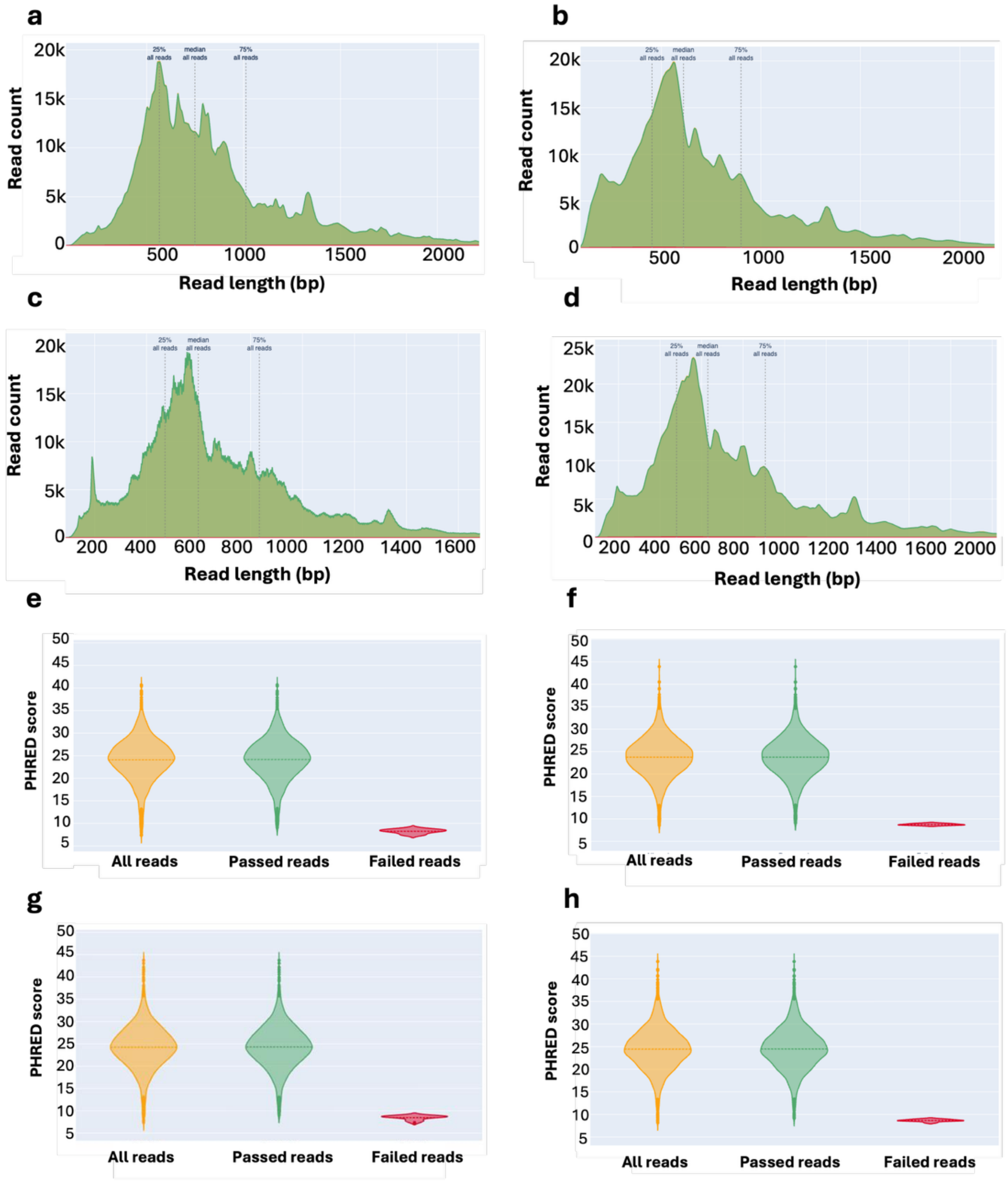


**Supplementary Figure 1: Representative Read Length and Quality Distributions. a**, Representative read length distribution for aged females. The x-axis indicates read length (bp), and the y-axis shows read counts ($\times 10^3$). Dotted lines denote read length quartiles, representing the distribution of read percentages across the sample. **b**, representative read length distribution for young females, displayed and interpreted as in the top panel. **c,** Representative read length distribution for aged males. The x-axis indicates read length (bp), and the y-axis shows read counts ($\times 10^3$). Dotted lines denote read length quartiles, representing the distribution of read percentages across the sample. **d,** Representative read length distribution for young males, displayed and interpreted as in the top panel. **e,** violin plot showing the representative Phred score distribution for aged samples. The y-axis indicates Phred score, and the x-axis denotes read type. **f**, Representative Phred score distribution for young female samples, displayed and interpreted as in the top panel. **g,** violin plot showing the representative Phred score distribution for aged males. The y-axis indicates Phred score, and the x-axis denotes read type. **h**, Representative Phred score distribution for young females, displayed and interpreted as in the top panel.

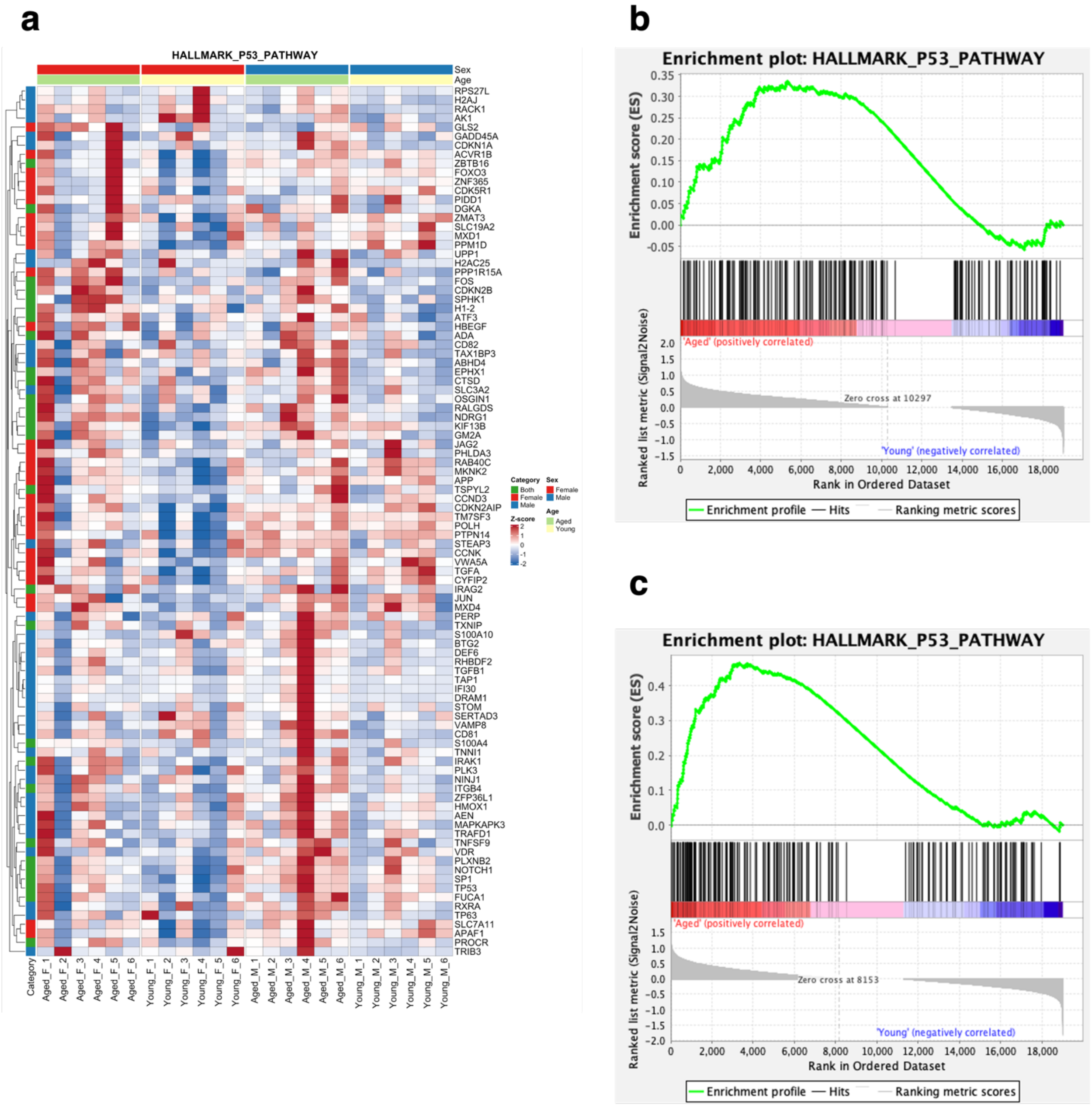


**Supplementary Figure 2: GSEA results for hallmark p53 pathway in aged vs young rhesus macaques. a,** Heatmap of normalized gene expression for genes in the ranked list of the HALLMARK_P53_PATHWAY from GSEA. Red indicates upregulation and blue indicates downregulation. Sex annotation: red, female; blue, male. Age annotation: yellow, young; green, aged. Category columns denote gene contributions to pathway enrichment: female-specific (red), male-specific (blue), or shared (green). **b,** Running ES plot for Aged_F versus Young_F groups. Genes are ranked by signal-to-noise, with Aged_F-enriched genes on the left (red) and Young_F on the right (blue). The ES profile (green) peaks in the leading-edge subset, indicating pathway upregulation in Aged_F. **c,** interpreted similarly to b. Running ES plot for Aged_M versus Young_M. The ES profile peaks toward Aged_M, indicating pathway upregulation relative to Young_M.

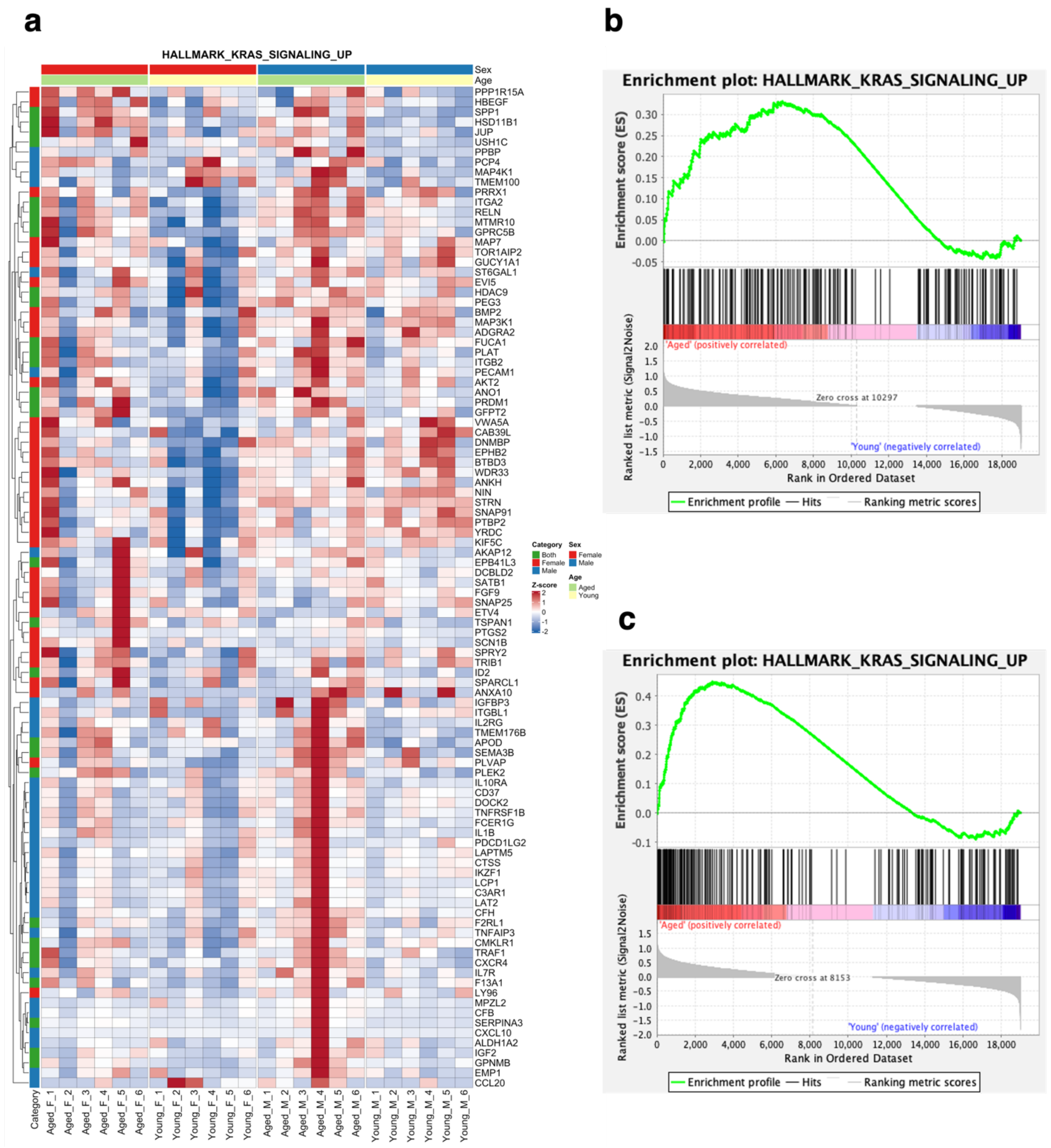


**Supplementary Figure 3: GSEA results for hallmark KRAS signaling pathway in aged vs young rhesus macaques. a,** Heatmap of normalized gene expression for genes in the ranked list of the HALLMARK_KRAS_SIGNALING_PATHWAY from GSEA. Red indicates upregulation and blue indicates downregulation. Sex annotation: red, female; blue, male. Age annotation: yellow, young; green, aged. Category columns denote gene contributions to pathway enrichment: female-specific (red), male-specific (blue). **b,** Running ES plot for Aged_F (aged females) versus Young_F (young females). Genes are ranked by signal-to-noise, with Aged_F-enriched genes on the left (red) and Young_F on the right (blue). The ES profile (green) peaks in the trailing-edge subset, indicating pathway upregulation in Young_F. **c,** interpreted similarly to b. Running ES plot for Aged_M (aged males) versus Young_M (young males). The ES profile peaks toward Aged_M, indicating pathway upregulation relative to Young_M.

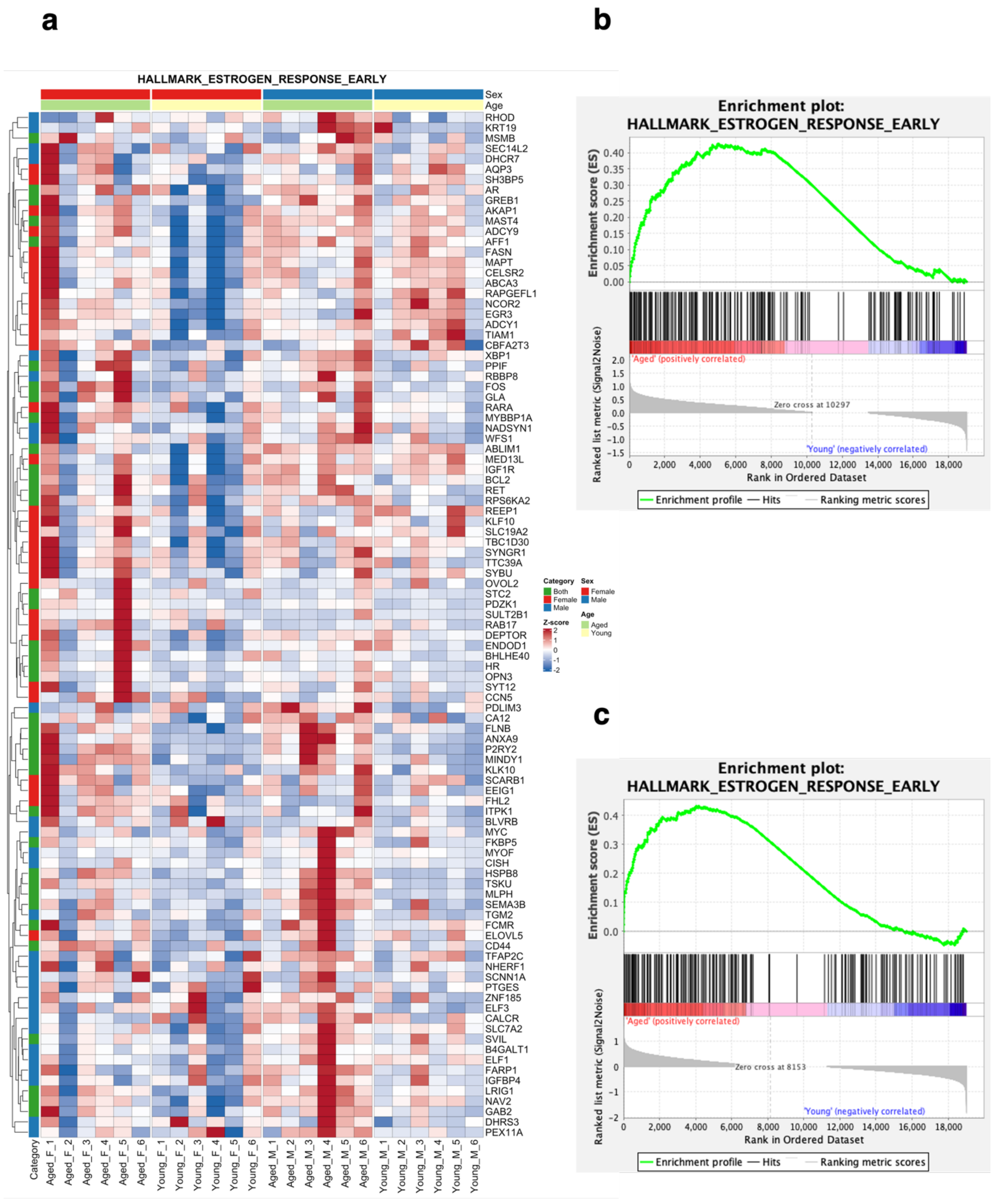
a
b
c
HALLMARK_ESTROGEN_RESPONSE_EARLY
Sex
Age
RHOD
KRT19
MSMB
SEC14L2
DHCR7
AQP3
SH3BP5
AR
GREB1
AKAP1
MAST4
ADCY9
AFF1
FASN
MAPT
CELSR2
ABCA3
RAPGEFL1
NCOR2
EGR3
ADCY1
TIAM1
CBFA2T3
XBP1
PPIF
RBBP8
FOS
GLA
RARA
MYBBP1A
NADSYN1
WFS1
ABLIM1
MED13L
IGF1R
BCL2
RET
RPS6KA2
REEP1
KLF10
SLC19A2
TBC1D30
SYNGR1
TTC39A
SYBU
OVOL2
STC2
PDZK1
SULT2B1
RAB17
DEPTOR
ENDOD1
BHLHE40
HR
OPN3
SYT12
CCN5
PDLIM3
CA12
FLNB
ANXA9
P2RY2
MINDY1
KLK10
SCARB1
EEIG1
FHL2
ITPK1
BLVRB
MYC
FKBP5
MYOF
CISH
HSPB8
TSKU
MLPH
SEMA3B
TGM2
FCMR
ELOVL5
CD44
TFAP2C
NHERF1
SCNN1A
PTGES
ZNF185
ELF3
CALCR
SLC7A2
SVIL
B4GALT1
ELF1
FARP1
IGFBP4
LRIG1
NAV2
GAB2
DHRS3
PEX11A
Category
Aged_F_1
Aged_F_2
Aged_F_3
Aged_F_4
Aged_F_5
Aged_F_6
Young_F_1
Young_F_2
Young_F_3
Young_F_4
Young_F_5
Young_F_6
Aged_M_1
Aged_M_2
Aged_M_3
Aged_M_4
Aged_M_5
Aged_M_6
Young_M_1
Young_M_2
Young_M_3
Young_M_4
Young_M_5
Young_M_6
Category
Both
Female
Male
Sex
Female
Male
Age
Aged
Young
Z-score
2
1
0
-1
-2
Enrichment plot:
HALLMARK_ESTROGEN_RESPONSE_EARLY
Enrichment score (ES)
'Aged' (positively correlated)
Zero cross at 10297
'Young' (negatively correlated)
Ranked list metric (Signal2Noise)
Rank in Ordered Dataset
Enrichment profile
Hits
Ranking metric scores
Enrichment plot:
HALLMARK_ESTROGEN_RESPONSE_EARLY
Enrichment score (ES)
'Aged' (positively correlated)
Zero cross at 8153
'Young' (negatively correlated)
Ranked list metric (Signal2Noise)
Rank in Ordered Dataset
Enrichment profile
Hits
Ranking metric scores

**Supplementary Figure 4: GSEA results for hallmark early estrogen response pathway in aged vs young rhesus macaques. a,** Heatmap of normalized gene expression for genes in the ranked list of the HALLMARK_ESTROGEN_RESPONSE_LATE from GSEA. Red indicates upregulation and blue indicates downregulation. Sex annotation: red, female; blue, male. Age annotation: yellow, young; green, aged. Category columns denote gene contributions to pathway enrichment: female-specific (red), male-specific (blue). **b,** Running ES plot for Aged_F (aged females) versus Young_F (young females). Genes are ranked by signal-to-noise, with Aged_F-enriched genes on the left (red) and Young_F on the right (blue). The ES profile (green) peaks in the trailing-edge subset, indicating positive gene set enrichment in Young_F and negative gene set enrichment in Aged_F. **c,** interpreted similarly to b. Running ES plot for Aged_M (aged males) versus Young_M (young males). The ES profile peaks toward Aged_M, indicating pathway upregulation relative to Young_M.

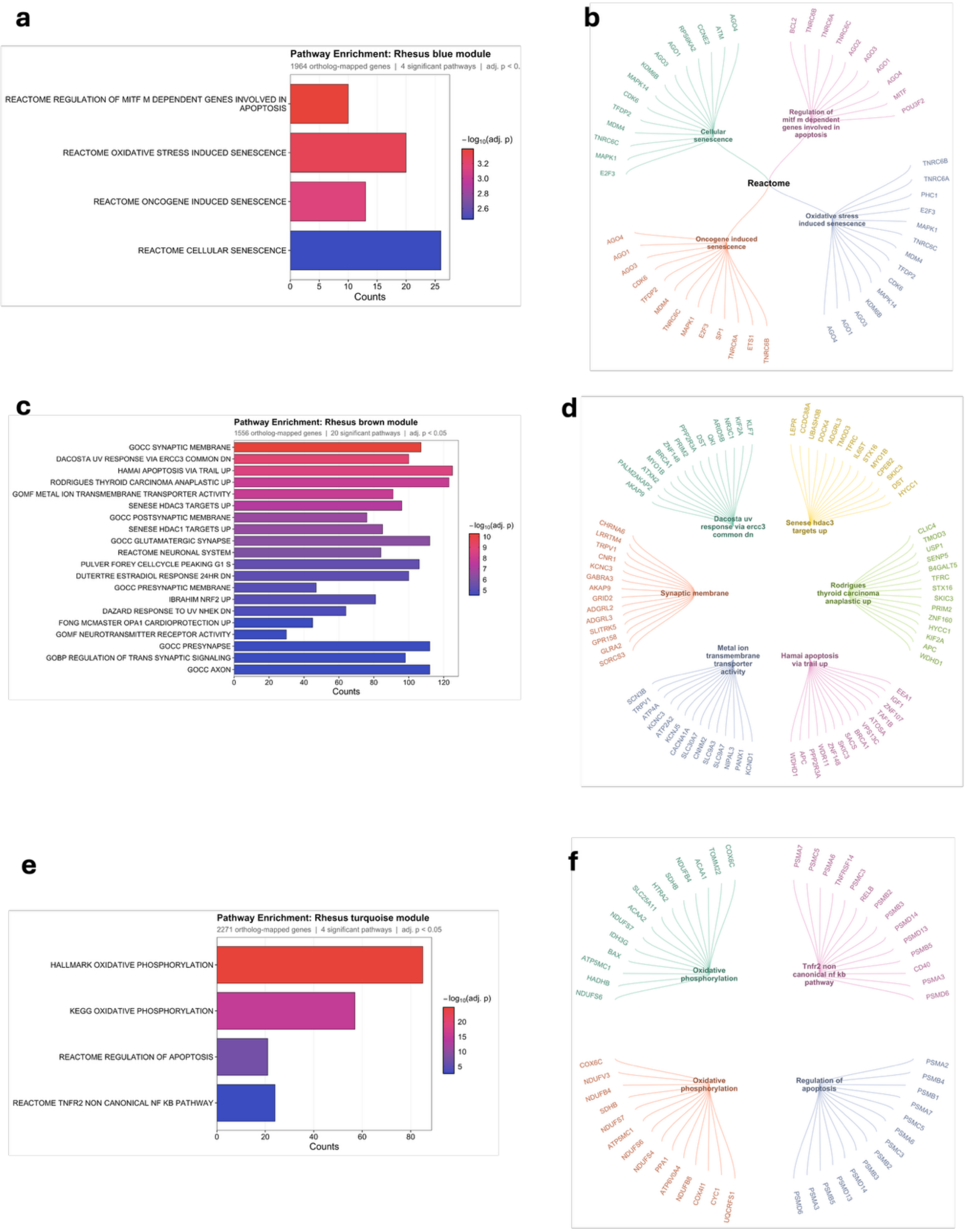

a
Pathway Enrichment: Rhesus blue module
1964 ortholog-mapped genes | 4 significant pathways | adj. p < 0.
REACTOME REGULATION OF MITF M DEPENDENT GENES INVOLVED IN APOPTOSIS
REACTOME OXIDATIVE STRESS INDUCED SENESCENCE
REACTOME ONCOGENE INDUCED SENESCENCE
REACTOME CELLULAR SENESCENCE
Counts
−log10(adj. p)
b
Reactome
Cellular senescence
Regulation of mitf m dependent genes involved in apoptosis
Oxidative stress induced senescence
Oncogene induced senescence
c
Pathway Enrichment: Rhesus brown module
1556 ortholog-mapped genes | 20 significant pathways | adj. p < 0.05
GOCC SYNAPTIC MEMBRANE
DACOSTA UV RESPONSE VIA ERCC3 COMMON DN
HAMAI APOPTOSIS VIA TRAIL UP
RODRIGUES THYROID CARCINOMA ANAPLASTIC UP
GOMF METAL ION TRANSMEMBRANE TRANSPORTER ACTIVITY
SENESE HDAC3 TARGETS UP
GOCC POSTSYNAPTIC MEMBRANE
SENESE HDAC1 TARGETS UP
GOCC GLUTAMATERGIC SYNAPSE
REACTOME NEURONAL SYSTEM
PULVER FOREY CELLCYCLE PEAKING G1 S
DUTERTRE ESTRADIOL RESPONSE 24HR DN
GOCC PRESYNAPTIC MEMBRANE
IBRAHIM NRF2 UP
DAZARD RESPONSE TO UV NHEK DN
FONG MCMASTER OPA1 CARDIOPROTECTION UP
GOMF NEUROTRANSMITTER RECEPTOR ACTIVITY
GOCC PRESYNAPSE
GOBP REGULATION OF TRANS SYNAPTIC SIGNALING
GOCC AXON
Counts
−log10(adj. p)
d
Dacosta uv response via ercc3 common dn
Senese hdac3 targets up
Synaptic membrane
Rodrigues thyroid carcinoma anaplastic up
Metal ion transmembrane transporter activity
Hamai apoptosis via trail up
e
Pathway Enrichment: Rhesus turquoise module
2271 ortholog-mapped genes | 4 significant pathways | adj. p < 0.05
HALLMARK OXIDATIVE PHOSPHORYLATION
KEGG OXIDATIVE PHOSPHORYLATION
REACTOME REGULATION OF APOPTOSIS
REACTOME TNFR2 NON CANONICAL NF KB PATHWAY
Counts
−log10(adj. p)
f
Oxidative phosphorylation
Tnfr2 non canonical nf kb pathway
Oxidative phosphorylation
Regulation of apoptosis

**Supplementary Figure 5: Pathway enrichment of rhesus macaque co-expression modules.**
**a-b,** Pathway enrichment and gene–pathway association networks for the rhesus blue module. **a,** Bar plot showing significantly enriched Reactome, KEGG, and Hallmark pathways in the blue module, with bar length indicating the number of module genes assigned to each pathway and color indicating enrichment significance ($-\log_{10}$ adjusted *P* value). Only pathways meeting an adjusted P-value <0.05 threshold are shown. **b,** Gene–pathway networks cnetplot depicting the genes contributing to the enriched pathways within the blue module, with each color denoting a distinct pathway. **c-d** Pathway enrichment and gene–pathway association networks for the rhesus brown module. **c,** Bar plot showing significantly enriched Reactome, KEGG, and Hallmark pathways in the brown module interpreted similarly to a, with bar length indicating the number of module genes assigned to each pathway and color indicating enrichment significance ($-\log_{10}$ adjusted *P* value). Only pathways meeting an adjusted P-value <0.05 threshold are shown. **d,** Gene–pathway networks cnetplot depicting the genes contributing to the enriched pathways within the brown module interpreted similarly to b, with each color denoting a distinct pathway. **e-f,** Pathway enrichment and gene–pathway association networks for the rhesus turquoise module. **e,** Bar plot showing significantly enriched Reactome, KEGG, and Hallmark pathways in the turquoise module interpreted similarly to a and c. **f,** Gene–pathway networks cnetplot depicting the genes contributing to the enriched pathways within the turquoise module interpreted similarly to b and d.

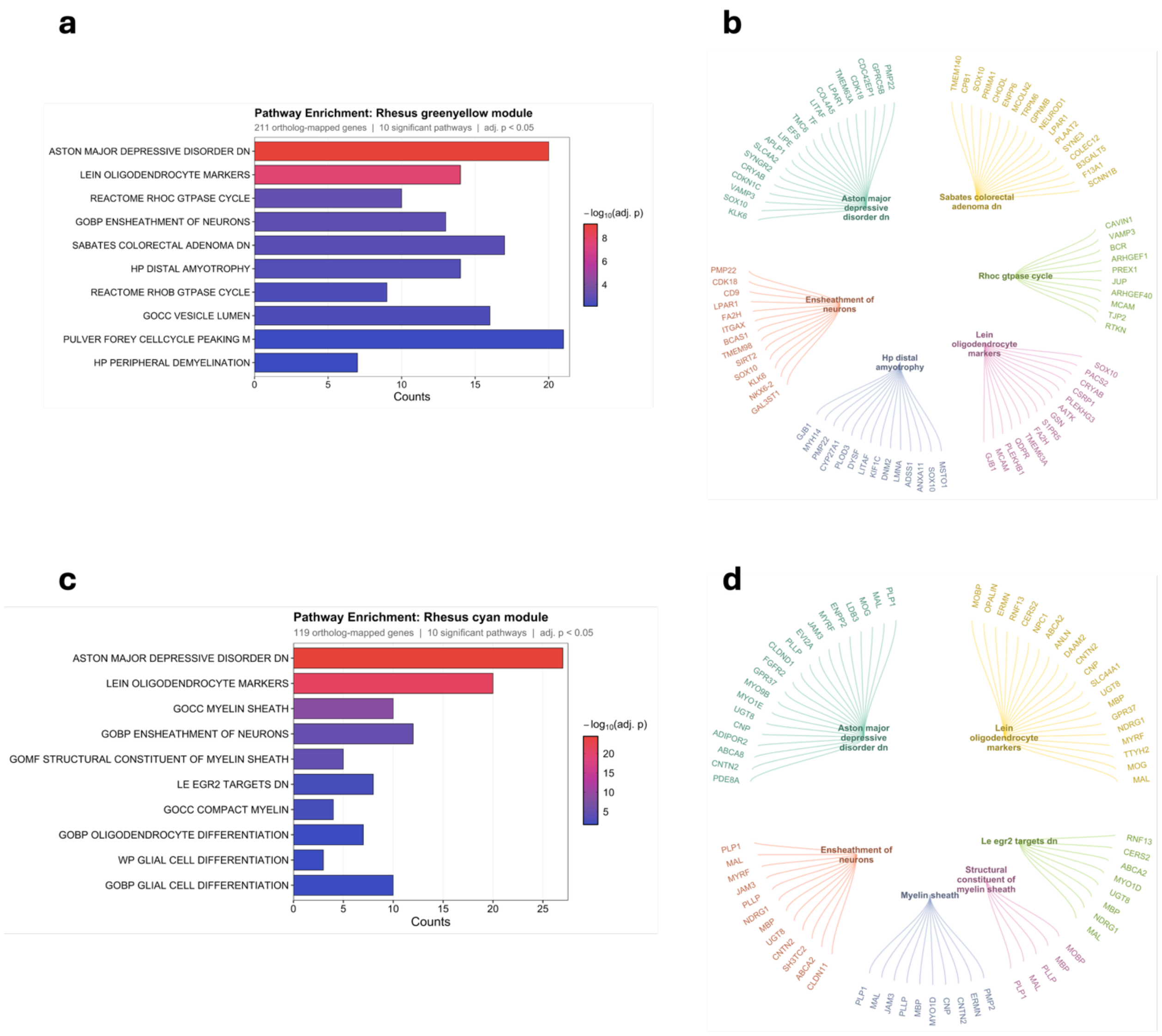


**Supplementary Figure 6: Pathway enrichment of male rhesus macaque co-expression modules.**
**a-b,** Pathway enrichment and gene–pathway association networks for the rhesus green-yellow module. **a,** Bar plot showing significantly enriched Reactome, KEGG, and Hallmark pathways in the green-yellow module, with bar length indicating the number of module genes assigned to each pathway and color indicating enrichment significance ($-\log_{10}$ adjusted *P* value). Only pathways meeting an adjusted P-value <0.05 threshold are shown. **b,** Gene–pathway networks cnetplot depicting the genes contributing to the enriched pathways within the green-yellow module, with each color denoting a distinct pathway. **c-d** Pathway enrichment and gene–pathway association networks for the rhesus cyan module. **c,** Bar plot showing significantly enriched Reactome, KEGG, and Hallmark pathways in the cyan module interpreted similarly to a, with bar length indicating the number of module genes assigned to each pathway and color indicating enrichment significance ($-\log_{10}$ adjusted *P* value). Only pathways meeting an adjusted P-value <0.05 threshold are shown. **d,** Gene–pathway networks cnetplot depicting the genes contributing to the enriched pathways within the cyan module interpreted similarly to b, with each color denoting a distinct pathway

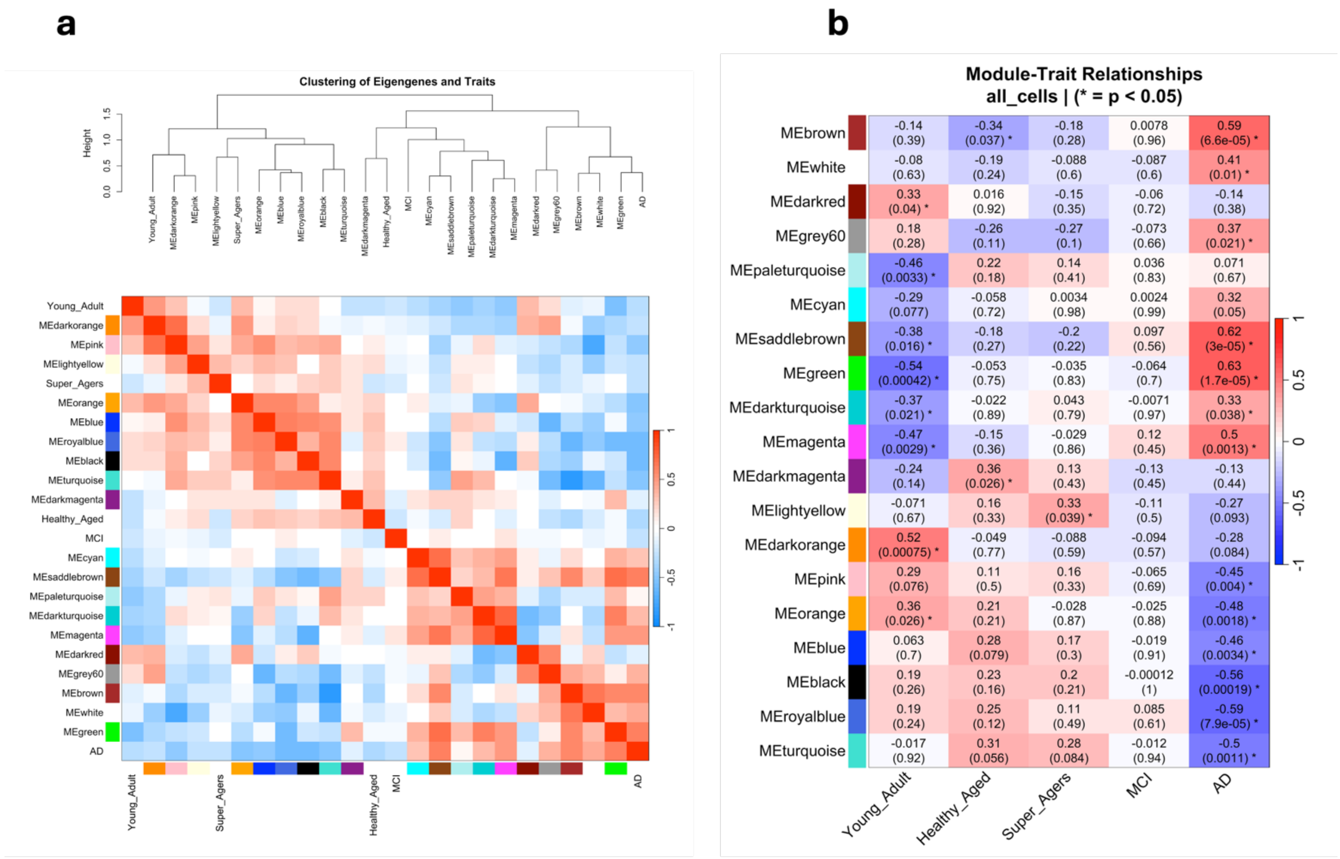


**Supplementary Figure 7: Human hippocampal WGCNA network structure and module–trait associations. a,** Clustering of significant module eigengenes and traits with the corresponding eigengene adjacency heat map. The dendrogram summarizes eigengene similarity, and the adjacent heat map depicts pairwise eigengene correlations, where warm and cool colors indicate positively and negatively correlated eigengenes, respectively. **b,** Heat map of Pearson correlations between module eigengenes and age–sex traits, highlighting modules significantly associated with young adults, healthy aged, super agers, mild cognitive impairment (MCI), and Alzheimer's Disease (AD). The color scale denotes correlation strength and sign (red, positive; blue, negative), and each cell reports the correlation coefficient with the corresponding P value in parentheses; asterisks indicate significant associations ($P < 0.05$).

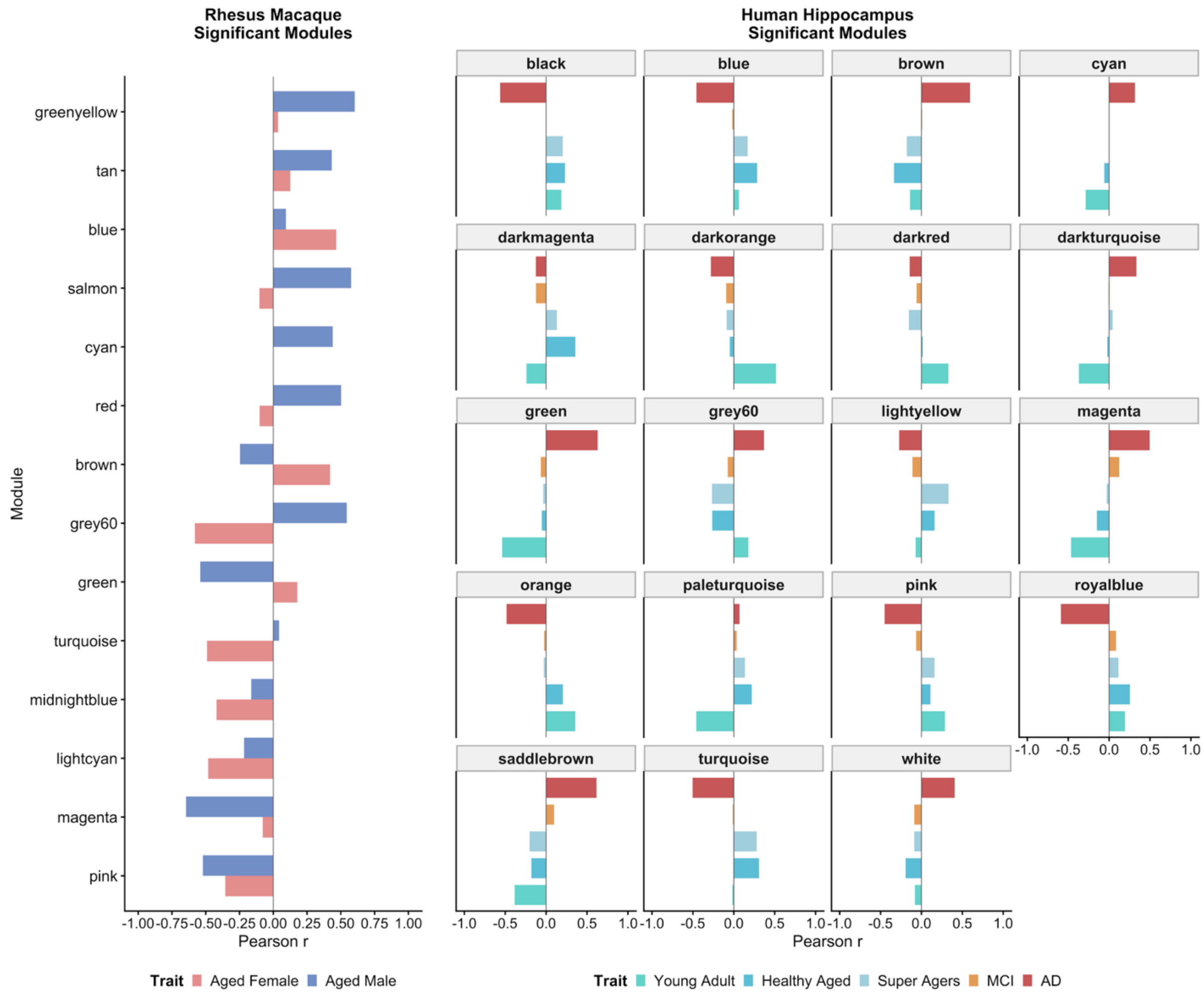


**Supplementary Figure 8: Significant WGCNA modules and their associations with aging and Alzheimer's disease traits in rhesus macaques and humans.** Pearson correlation coefficients between significant weighted gene co-expression network analysis (WGCNA) module eigengenes and phenotypic traits are shown for rhesus macaque (left) and human hippocampal (right) datasets. For the rhesus macaque network, significant module eigengenes were correlated with aged female and aged male groups. For the human hippocampal network, significant module eigengenes were correlated with young adults, healthy aged individuals, Super Agers, individuals with mild cognitive impairment (MCI), and individuals with Alzheimer's disease (AD). Positive and negative Pearson correlation coefficients are represented by bars extending to the right and left of zero, respectively. Only modules significantly associated with at least one phenotypic trait (FDR-adjusted $P < 0.05$) were retained for downstream cross-species network comparison.

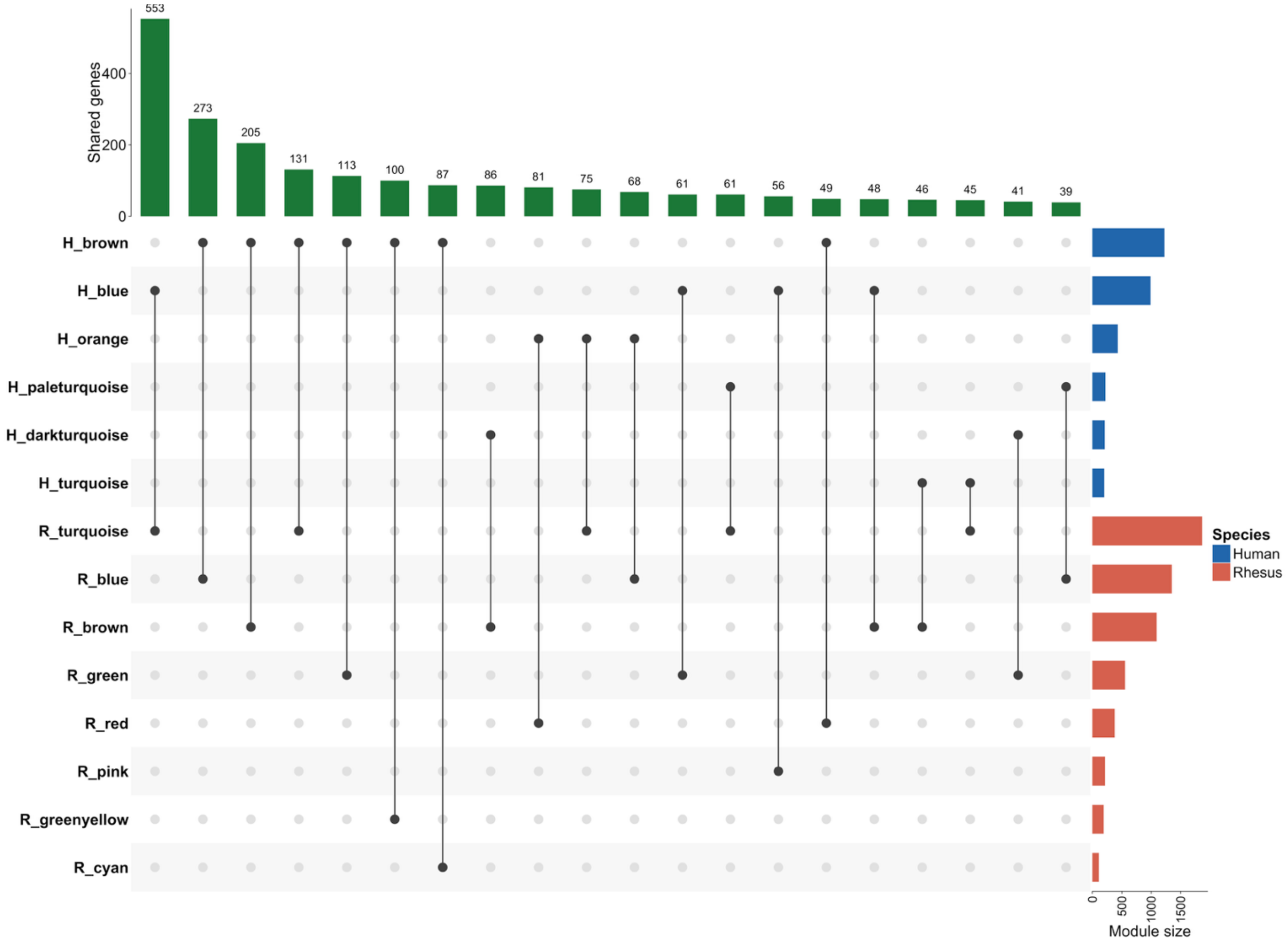


**Supplementary Figure 9: UpSet plot of overlapping genes between rhesus macaque and human WGCNA modules.** UpSet plot illustrating the overlap of genes between rhesus macaque (orange) and human (blue) weighted gene co-expression network analysis (WGCNA) modules. Horizontal bar plots indicate the total number of genes within each rhesus and human module, while the vertical bar plot represents the number of genes shared between module pairs. Connected dots below the vertical bars identify the corresponding rhesus and human modules with shared genes.

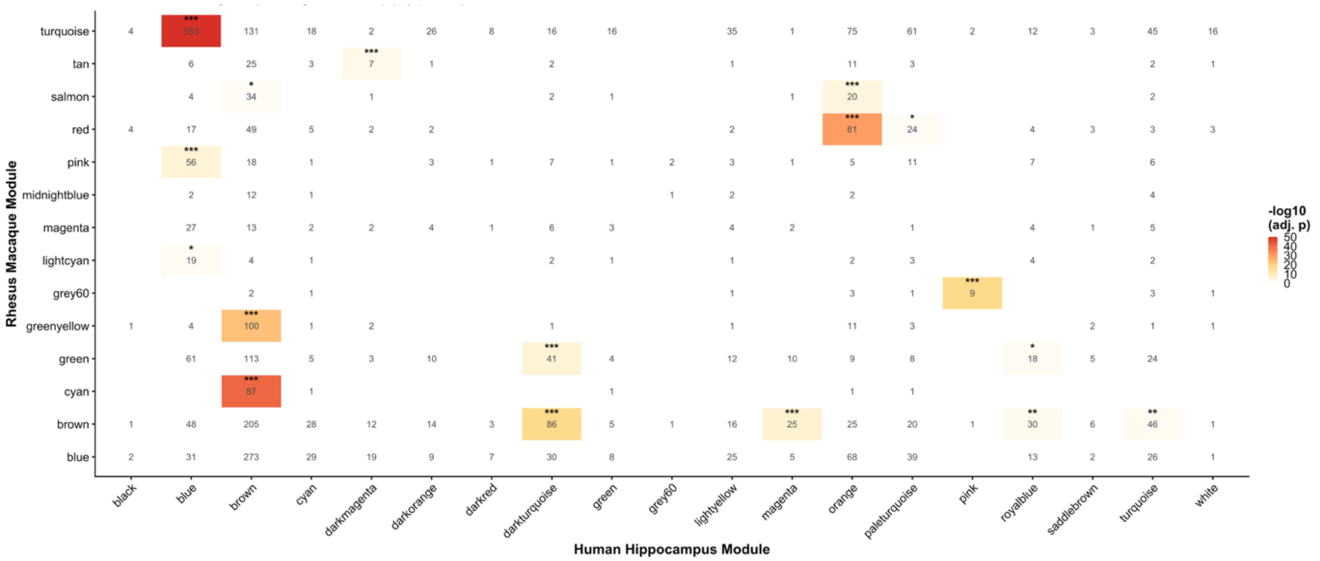


**Supplementary Figure 10: Pairwise gene overlap matrix between significant rhesus macaque and human hippocampal co-expression modules.**

Heatmap showing the number of shared ortholog-mapped genes between significant rhesus macaque aging modules (rows) and significant human hippocampal co-expression modules (columns). Values within cells indicate the number of overlapping ortholog-mapped genes. Cell color represents the statistical significance of overlap, expressed as −log10 Benjamini–Hochberg-adjusted P value from a hypergeometric test evaluating whether the observed overlap exceeds that expected by chance given module size and the ortholog-mapped gene universe. White cells indicate non-significant overlap; increasing color intensity indicates greater significance. Statistically significant overlaps are marked with asterisks (adjusted P < 0.05). Seventeen module pairs exhibited significant cross-species overlap, with the strongest associations observed for rhesus turquoise–human blue (553 shared genes; adjusted P = 2.20 × 10−132), rhesus cyan–human brown (87 shared genes; adjusted P = 2.25 × 10−41), rhesus red–human orange (81 shared genes; adjusted P = 6.63 × 10−31), and rhesus grey60–human pink (9 shared genes; adjusted P = 1.38 × 10−18). Empty cells indicate no shared genes between module pairs.

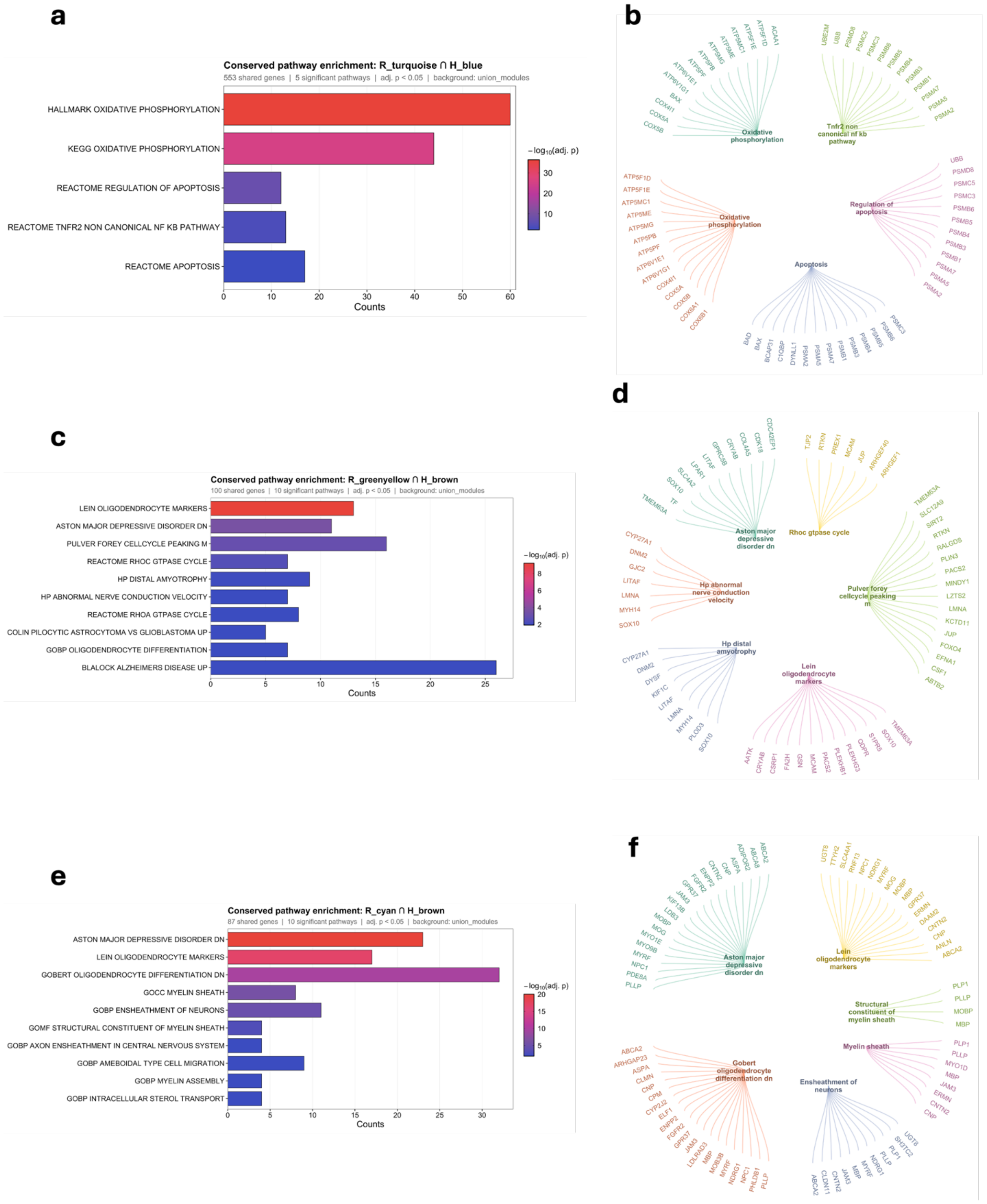
a
Conserved pathway enrichment: R_turquoise ∩ H_blue
553 shared genes | 5 significant pathways | adj. p < 0.05 | background: union_modules
HALLMARK OXIDATIVE PHOSPHORYLATION
KEGG OXIDATIVE PHOSPHORYLATION
REACTOME REGULATION OF APOPTOSIS
REACTOME TNFR2 NON CANONICAL NF KB PATHWAY
REACTOME APOPTOSIS
Counts
−log10(adj. p)
b
Oxidative phosphorylation
Tnfr2 non canonical nf kb pathway
Regulation of apoptosis
Apoptosis
c
Conserved pathway enrichment: R_greenyellow ∩ H_brown
100 shared genes | 10 significant pathways | adj. p < 0.05 | background: union_modules
LEIN OLIGODENDROCYTE MARKERS
ASTON MAJOR DEPRESSIVE DISORDER DN
PULVER FOREY CELLCYCLE PEAKING M
REACTOME RHOC GTPASE CYCLE
HP DISTAL AMYOTROPHY
HP ABNORMAL NERVE CONDUCTION VELOCITY
REACTOME RHOA GTPASE CYCLE
COLIN PILOCYTIC ASTROCYTOMA VS GLIOBLASTOMA UP
GOBP OLIGODENDROCYTE DIFFERENTIATION
BLALOCK ALZHEIMERS DISEASE UP
Counts
−log10(adj. p)
d
Aston major depressive disorder dn
Rhoc gtpase cycle
Hp abnormal nerve conduction velocity
Pulver forey cellcycle peaking m
Hp distal amyotrophy
Lein oligodendrocyte markers
e
Conserved pathway enrichment: R_cyan ∩ H_brown
87 shared genes | 10 significant pathways | adj. p < 0.05 | background: union_modules
ASTON MAJOR DEPRESSIVE DISORDER DN
LEIN OLIGODENDROCYTE MARKERS
GOBERT OLIGODENDROCYTE DIFFERENTIATION DN
GOCC MYELIN SHEATH
GOBP ENSHEATHMENT OF NEURONS
GOMF STRUCTURAL CONSTITUENT OF MYELIN SHEATH
GOBP AXON ENSHEATHMENT IN CENTRAL NERVOUS SYSTEM
GOBP AMEBOIDAL TYPE CELL MIGRATION
GOBP MYELIN ASSEMBLY
GOBP INTRACELLULAR STEROL TRANSPORT
Counts
−log10(adj. p)
f
Aston major depressive disorder dn
Lein oligodendrocyte markers
Structural constituent of myelin sheath
Myelin sheath
Gobert oligodendrocyte differentiation dn
Ensheathment of neurons

**Supplementary Figure 11: Pathway enrichment of rhesus macaque and human shared genes.**
**a-b,** Pathway enrichment and gene–pathway association networks of genes conserved between rhesus turquoise and human blue. **a,** Bar plot showing significantly enriched Reactome, KEGG, and Hallmark pathways of genes conserved between rhesus turquoise and human blue, with bar length indicating the number of module genes assigned to each pathway and color indicating enrichment significance ($-\log_{10}$ adjusted *P* value). Only pathways meeting an adjusted P-value <0.05 threshold are shown. **b,** Gene–pathway networks cnetplot depicting the genes contributing to the enriched pathways of genes conserved between rhesus turquoise and human blue, with each color denoting a distinct pathway. **c-d** Pathway enrichment and gene–pathway association networks of genes conserved between rhesus green-yellow and human brown. **c,** Bar plot showing significantly enriched Reactome, KEGG, and Hallmark pathways of genes conserved between rhesus green-yellow and human brown interpreted similarly to a, with bar length indicating the number of module genes assigned to each pathway and color indicating enrichment significance ($-\log_{10}$ adjusted *P* value). Only pathways meeting an adjusted P-value <0.05 threshold are shown. **d,** Gene–pathway networks cnetplot depicting the genes contributing to the enriched pathways within of genes conserved between rhesus green-yellow and human brown interpreted similarly to b, with each color denoting a distinct pathway. **e-f,** Pathway enrichment and gene–pathway association networks of genes conserved between rhesus cyan and human brown. **e,** Bar plot showing significantly enriched Reactome, KEGG, and Hallmark pathways of genes conserved between rhesus cyan and human brown interpreted similarly to a and c. **f,** Gene–pathway networks cnetplot depicting the genes contributing to the enriched pathways conserved between rhesus cyan and human brown interpreted similarly to b and d.

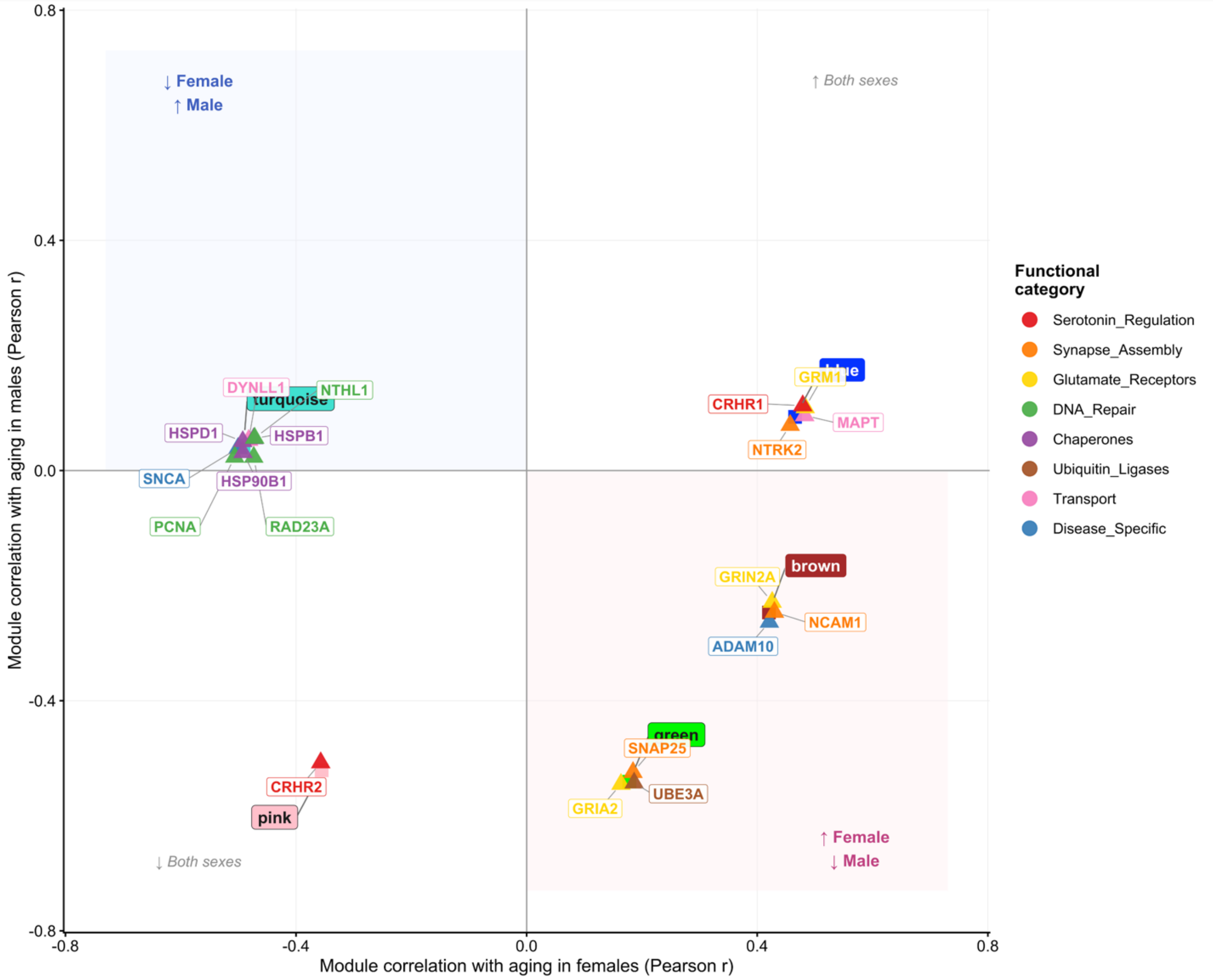


**Supplementary Figure 12: Sex-Differential Aging Effect of Estrogen-Regulated Genes in Significant Rhesus Hippocampal Co-Expression Modules.** Quadrant dot plot showing the sex-differential associations between aging and 13 estrogen-regulated genes identified within significant rhesus macaque hippocampal WGCNA co-expression modules. Each dot represents an individual gene. The x-axis denotes the Pearson correlation between module eigengene expression and age in females (Aged_Female), whereas the y-axis denotes the corresponding correlation in males (Aged_Male). Because module–trait correlations are calculated at the module level, genes assigned to the same module share the same underlying x–y coordinates and are displayed with slight random jitter for visual separation. Module positions are indicated by colored squares and labelled callouts, with connecting lines linking each gene to its corresponding module anchor. Gene colors denote functional categories defined in the estrogen-regulated gene reference list. Shaded quadrants indicate concordant sex-specific aging directions, with the upper-left and lower-right quadrants representing opposite aging associations between females and males; annotations indicate the direction of the aging association in each sex.
